# Cohesin constrains histone modification-driven chromatin dynamics

**DOI:** 10.64898/2025.12.13.694149

**Authors:** Takeshi Fujino, Yotaro Ochi, Susumu Goyama, Yuko Shimosato, Shiori Shikata, Ayana Kon, Takuto Mori, Toyonori Sakata, Hiroaki Honda, Haruhiko Koseki, Manabu Nakayama, Katsuhiko Shirahige, Satoru Miyano, Seishi Ogawa, Toshio Kitamura

**Author notes:** Co-correspondance (S.O.), (T.K.). These authors contributed equally (Takeshi Fujino, Yotaro Ochi). These authors contributed equally.

## Abstract

Gene expressions are regulated by an interplay between epigenetics and spatial genome organization, the deregulation of which has been implicated in the development of cancers, including myeloid neoplasms. However, it is unclear how they coordinately contribute to normal and malignant hematopoiesis. Here, we show that simultaneous dysregulations of histone modifications and chromatin structures caused by mutations of the epigenetic modulator *Asxl1* and cohesin subunit *Stag2* cooperatively induce ectopic interactions between polycomb-regulated promoters and active enhancers, leading to the aberrant upregulation of hematopoietic stem cell related genes and development of myelodysplastic syndromes (MDS). De-repression of polycomb-regulated genes induces their translocation to the transcriptionally active loci, where active promoters and enhancers are assembled, in the absence of Stag2-mediated chromatin organization. Our findings revealed that cohesin counteracts histone modification-driven chromatin conformations, manifesting the coordinate roles of histone modifiers and cohesin in regulating genome architectures and gene expressions to prevent malignant transformation.

## INTRODUCTION

Multi-modal regulatory mechanisms orchestrate gene expression programs to determine cell fate and cellular identity during cell differentiation and development. Here, extracellular and intracellular cues are transmitted through signaling pathways to converge on a series of intranuclear processes, changing epigenetic modifications, chromosomal conformations, and gene expressions ^1–4^. Genome-wide chromosome conformation capture (Hi-C) experiments have revealed that chromosomes are partitioned into contact domains with distinctive patterns of histone modifications ^5,6^. Locally, each genomic domain organizes loop-like structures to be spatially segregated into topologically associating domains (TADs) ^7,8^. TAD boundaries are demarcated by architectural proteins CCCTC-binding factor (CTCF) and cohesin, where CTCF stalls cohesin to halt loop extrusion according to the loop extrusion model ^9–11^. Depletion of cohesin reduces TAD organization and increases the compartmentalization of chromatin domains with similar histone modifications, suggesting that cohesin-mediated TAD formation competes with epigenetics-associated compartmentalization ^12,13^.

Each TAD functions as a transcriptional regulatory unit, where communication between promoters and enhancers is restricted within an insulated region ^14–16^. It has been proposed that multiple regulatory elements, including cohesin, mediator, and transcription factors, can form promoter-enhancer (P-E) interactions ^17–19^. Notably, a recent study showed that polycomb-regulated genes can form self-interactions upon removal of cohesin ^20^. Moreover, interactions between promoters and enhancers with active histone marks are increased in the absence of cohesin ^21^. These findings and the aforementioned epigenetics-associated compartmentalization illustrate the cooperative role of epigenetic modifications and cohesin in the regulation of the chromatin architecture and gene expressions.

In humans, the epigenetic modulator *ASXL1* and the cohesin subunit *STAG2* are recurrently mutated in patients with myeloid neoplasms, including MDS and AML ^22–24^. Mutations in *ASXL1* are frameshift or nonsense mutations in the last exon and lead to generation of the C-terminally truncated form of ASXL1 ^25–28^. It has been shown that the C-terminally truncated form of mutant Asxl1 (Asxl1-MT) de-represses polycomb-regulated genes by reducing H3K27me3 and H2AK119Ub levels ^29–31^. Meanwhile, *STAG2* mutations are frameshift or nonsense mutations located across the coding regions, resulting in a loss of function ^22^. Previous studies have shown that *STAG2* loss alters higher-order genome structures, including P-E interactions, to dysregulate gene expressions ^21,23,32,33^. We previously revealed mutations in *ASXL1* and *STAG2* frequently coexist in MDS patients ^22,23^, indicating that dysregulation of histone modifications and cohesin-mediated genomic structures confers a synergistic impact on myeloid transformation. These findings prompted us to investigate the interplay between epigenetic dynamics and chromatin architectures in the homeostatic and disease states.

## RESULTS

### Mutations in *ASXL1* and *STAG2* frequently coexist in MDS patients

To delineate the mutational landscape of MDS patients, we analyzed clinical data on patients with MDS (N = 2,498) and related myeloid neoplasms (N = 549) in our cohort and publicly available database ^23^. *ASXL1* and *STAG2* mutations were detected in 524 (17.2%) and 208 (6.8%) patients, respectively, of which concomitant mutations in *ASXL1* and *STAG2* (hereinafter referred to as “double mutations”) were found in 103 patients, accounting for 19.7% of *ASXL1*-mutated cases and 49.5% of *STAG2*-mutated cases **(Figure 1A)**. We next examined the clinical features of MDS patients with or without mutations in *ASXL1* and *STAG2*. We observed a higher frequency of high-risk MDS (RAEB) in patients harboring double mutations compared with those with *ASXL1* or *STAG2* mutation alone or neither mutation **(Figure S1A)**. There were no significant differences in the age or gender between patients among the mutation groups **(Figure S1B and S1C)**. Double mutations and *STAG2* mutation alone were similarly associated with a higher score of the revised International Prognostic Scoring System (IPSS-R), fewer white blood cells, fewer platelets, and higher frequency of blast cells **(Figure S1D-F)**. The characteristic finding of patients with double mutations was a low hemoglobin level **(Figure S1E)**. Lastly, patients with double mutations had worse overall survival compared to those with single or no mutation in *ASXL1* and *STAG2* (Figure 1B).

**Figure 1.**
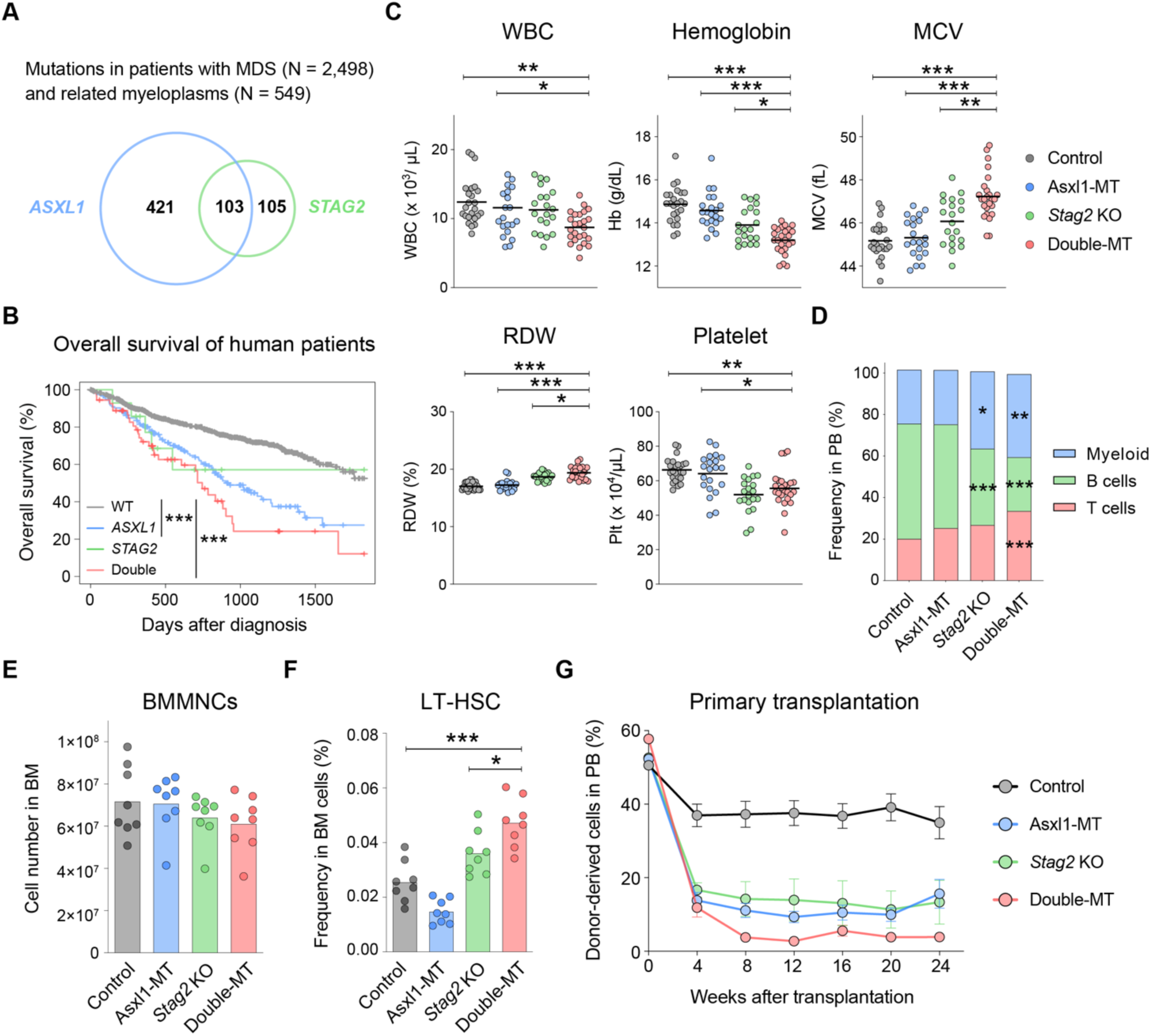
Synergistic impact of Asxl1-MT and *Stag2* loss on hematopoiesis. **(A)** Venn diagram showing the mutational overlap between *ASXL1* and *STAG2* in human patients with MDS (N = 2,498) and related myeloplasms (N = 549). **(B)** Kaplan-Meier curve of human MDS patients categorized by the presence of *ASXL1* or *STAG2* mutations. **(C)** Enumeration of white blood cells (WBCs), hemoglobin (Hb), mean corpuscular volume (MCV), red cell distribution width (RDW), and platelets (Plt) in peripheral blood from 12-week-old mice. N = 24 (Control), 21 (Asxl1-MT), 20 (*Stag2* KO), or 25 (double-MT) mice/group. Bar indicates mean value. **(D)** The frequencies of myeloid (CD11b^+^), B (B220^+^), and T (CD3^+^) cells in peripheral blood WBCs. N = 24 (Control), 21 (Asxl1-MT), 20 (*Stag2* KO), or 25 (double-MT) mice/group. Bar graphs indicate mean value. **(E)** The number of bone marrow mononuclear cells (BMMNCs). N = 8 mice/group. Bar graphs indicate mean value. **(F)** The frequency of long-term hematopoietic stem cells (LT-HSCs) in BMMNCs. N = 8 mice/group. Bar graphs indicate mean value. **(G)** 5 × 10^5^ BMMNCs isolated from control, ASXL1-MT, *Stag2* KO, or double-MT mice and 5 × 10^5^ wild-type BMMNCs were competitively transplanted into lethally irradiated recipient mice. The levels of donor chimerism in peripheral blood were analyzed at the indicated weeks after transplantation. N = 12 (Control), 15 (Asxl1-MT), 12 (*Stag2* KO), or 15 (double-MT) mice/group. Error bars indicate mean values ± S.E. Statistical significance was assessed by the log-ranked test **(B)** or one-way ANOVA with Tukey-Kramer’s post-hoc test **(C-G)**. *P ≤ 0.05, **P ≤ 0.01, ***P ≤ 0.001.

### Asxl1-MT and *Stag2* loss synergistically perturb hematopoiesis

To interrogate the mechanistic link between the epigenetics and chromatin architecture in hematopoiesis, we crossbred hematopoietic lineage-specific *Vav-Cre* conditional knockin (KI) mice expressing the C-terminally truncated form of ASXL1 (Asxl1-MT) and *Stag2* knockout (KO) mice to generate *Asxl1*/*Stag2* double-mutant (double-MT) mice. In accordance with previous reports ^23,31,34^, peripheral blood counts at 12 weeks of age were almost normal in Asxl1-MT KI mice, whereas *Stag2* KO mice showed anemia and thrombocytopenia. Double-MT mice also exhibited the same abnormal peripheral blood count as *Stag2* KO mice but were more anemic and had a decreased white blood cell count. Anemia in *Stag2* KO and double-MT mice was associated with elevated mean corpuscular volume (MCV) and red cell distribution width (RDW). These phenotypes were more prominent in double-MT mice compared with *Stag2* KO mice **(Figure 1C)**. In both *Stag2* KO and double-MT mice, the frequency of CD71^+^Ter119^+^ erythroid progenitors was decreased in the bone marrow but increased in the spleen **(Figure S2A)**, suggesting the presence of extramedullary hematopoiesis. *Stag2* KO and double-MT mice presented a skewed differentiation into myeloid lineage with a reduced fraction of B cells in the peripheral blood and bone marrow **(Figure 1D and S2B)**.

We next analyzed bone marrow hematopoietic stem and progenitor cell (HSPC) compartment. Asxl1-MT KI, *Stag2* KO, and double-MT mice showed normal bone marrow cellularity **(Figure 1E)**. Consistent with anemia and myeloid-skewed differentiation, there was a substantial increase in the granulocyte/macrophage progenitors (GMPs) and a decrease in the megakaryocyte/erythroid progenitors (MEPs) in double-MT mice **(Figure S2C).** There were no significant changes in the frequencies of Lin^-^Sca1^+^c-kit^+^ (LSK) cells, multipotent progenitors (MPPs; CD48^+^CD150^-^ LSK), or short-term HSCs (ST-HSCs; CD48^-^CD150^-^ LSK) in double-MT mice **(Figure S2D)**. However, as previously reported ^23,31,34,35^, Asxl1-MT decreased the frequency of long-term HSCs (LT-HSCs; CD48^-^CD150^+^ LSK), whereas *Stag2* loss increased it **(Figure 1F)**. Intriguingly, double-MT mice showed a highly increased LT-HSC fraction compared with control, Asxl1-MT KI mice and even *Stag2* KO mice. Although the increased LT-HSCs in double-MT mice suggested an expansion of the stem cell pool, they presented impaired repopulating potential accompanied by skewed differentiation into myeloid lineage **(Figure 1G, S2E, and S2F)**. Collectively, these findings demonstrate that concomitant mutations in *Asxl1* and *Stag2* synergistically promote expansion of LT-HSCs with impaired differentiation potential and perturb hematopoiesis.

### Concurrent mutations in *Asxl1* and *Stag2* lead to development of hematological malignancies

Similar to human patients, progressive macrocytic anemia was a conspicuous feature that discriminated double-MT mice from Asxl1-MT KI and *Stag2* KO mice during long-term follow-up **(Figure 2A, S3A, and S3B)**. Blood smears from double-MT mice revealed the presence of white blood cells with morphological abnormalities compatible with MDS **(Figure S3C)**. All double-MT mice eventually died of hematological neoplasms at the age of 6 to 18 months (**Figure 2B**). The most common cause of death was MDS (8 out of 12 mice: 67%) (**Figure 2C**), followed by AML (3 out of 12 mice: 25%) and B-ALL (1 out of 12 mice: 8.3%); neither Asxl1-MT KI nor *Stag2* KO mice died of such neoplasms **(Figure 2D)**. Importantly, mice that developed AML had exhibited dysplastic blood cells before the onset of AML, suggesting the presence of preceding MDS **(Figure S3D)**.

**Figure 2.**
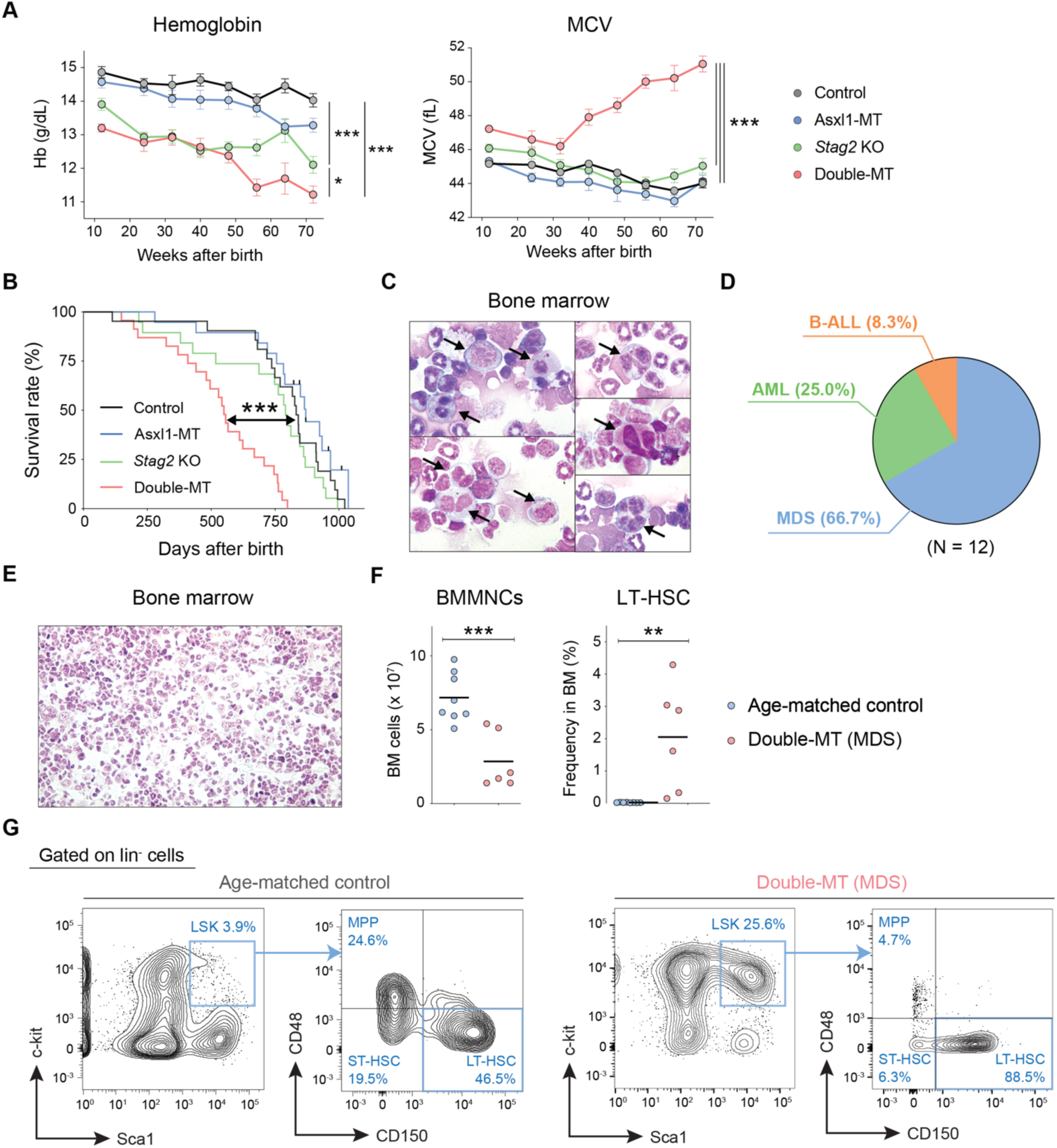
Concurrent mutations of *Asxl1* and *Stag2* induce development of MDS in mice. **(A)** Enumeration of Hb and MCV at the indicated weeks after birth. N = 24 (Control), 21 (Asxl1-MT), 20 (*Stag2* KO), or 25 (double-MT) mice/group. Error bars indicate mean values ± S.E. **(B)** Kaplan-Meier survival curves. N = 24 (Control), 21 (Asxl1-MT), 20 (*Stag2* KO), or 25 (double-MT) mice/group. **(C)** Microscopic images of Wright-Giemsa-stained bone marrow cells from double-MT mice died of MDS. **(D)** Pie chart showing the cause of death of double-MT mice (N = 12). **(E-G)** Analyses of bone marrow cells in moribund double-MT mice with MDS. **(E)** Hematoxylin-Eosin staining of bone marrow cells. **(F)** The number of bone marrow MNCs and LT-HSCs in double-MT mice and age-matched control mice. N = 8 (age-matched control) or 6 (double-MT). Bars indicate mean values. **(G)** Representative flow cytometry plots of bone marrow lin^-^ cells from age-matched control (left panel) and double-MT mice (right panel) with MDS. Statistical significance was assessed by one-way ANOVA with Tukey-Kramer’s post-hoc test **(A)**, log-ranked test **(B)**, or two-tailed Student’s t-test **(F)**. *P ≤ 0.05, **P ≤ 0.01, ***P ≤ 0.001.

We attempted to delineate pathogenesis of MDS and AML in double-MT mice. The mice with MDS presented hypocellular bone marrow and abnormal expansion of LT-HSCs **(Figure 2E-G)**, whereas there was a monotonous proliferation of GMPs or CD34^-^CD16/32^+^ Lin^-^c^-^kit^+^ myeloid progenitors in mice with AML, reminiscent of leukemic GMPs (L-GMPs) ^36^ **(Figure S3E)**. These observations imply that the abnormal proliferation of myeloid progenitors and HSCs are characteristic of AML and MDS, respectively^37^. Taken together, these data suggest that Asxl1-MT collaborates with *Stag2* loss to expand dysfunctional HSCs, resulting in ineffective hematopoiesis and development of MDS.

### Asxl1-MT and *Stag2* loss synergistically dysregulate gene expression

To evaluate the synergistic effect of *Asxl1* and *Stag2* mutations on gene expressions, we performed RNA sequencing (RNA-seq) analysis of bone marrow LSK cells from young mice (12 weeks). A principal component analysis (PCA) distinctively clustered LSK cells from control, Asxl1-MT KI, *Stag2* KO, and double-MT mice (**Figure 3A**). LSK cells of double-MT mice had a higher number of differentially expressed genes (vs. control) compared to those of Asxl1-MT KI or *Stag2* KO mice (**Figure 3B**). A pathway analysis of double-MT-specific deferentially expressed genes (**Figure S4A**; defined as genes with significant expression change in double-MT mice, but not in Asxl1-MT KI or *Stag2* KO mice) revealed the upregulation of HSC signature genes and downregulation of differentiation-related genes (**Figure 3C and S4B**), consistent with abnormal expansion of HSCs and ineffective hematopoiesis in double-MT mice.

**Figure 3.**
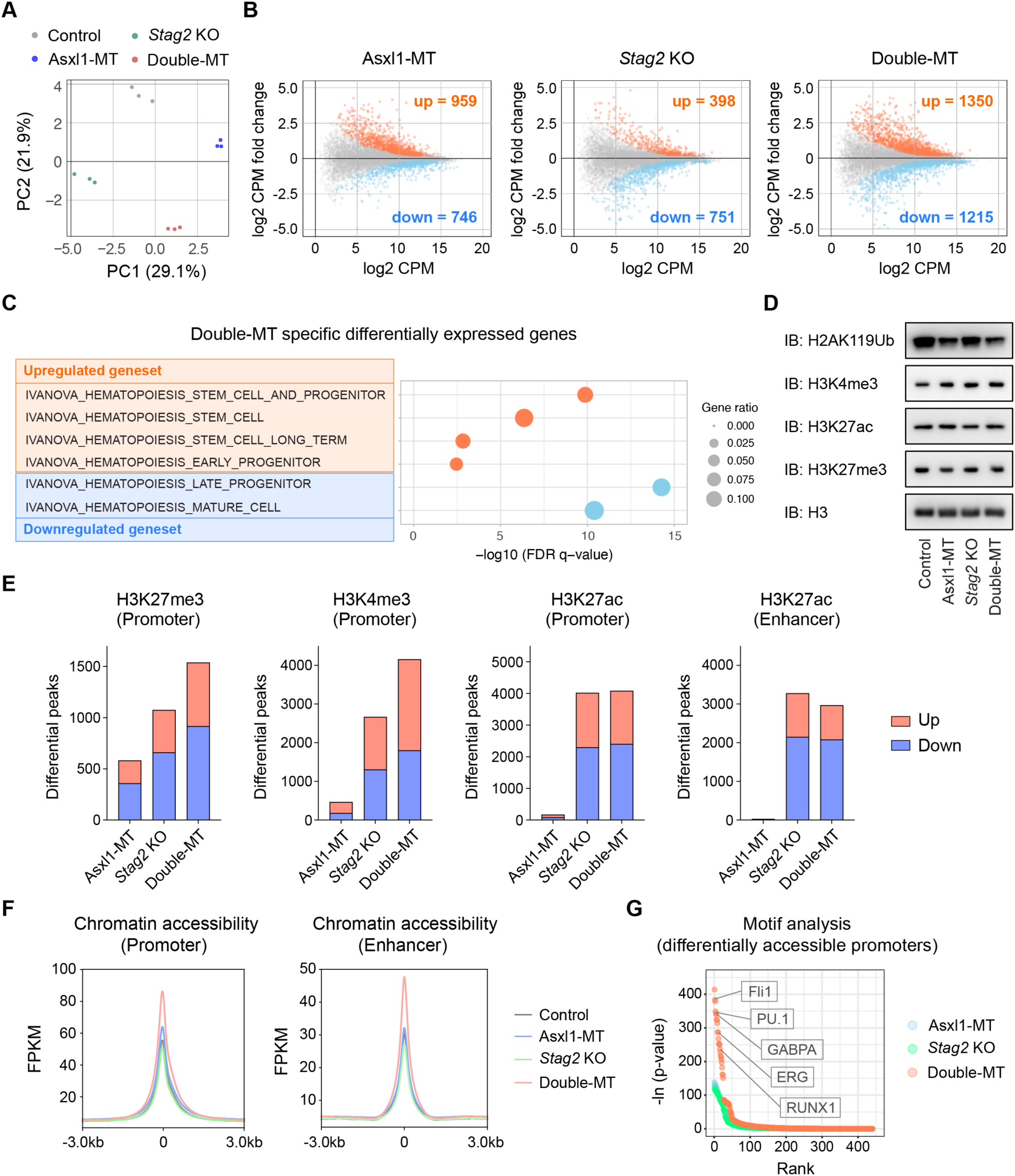
Asxl1-MT and *Stag2* loss synergistically disrupt gene expressions accompanied by changes in histone modifications and chromatin accessibility. (A, B) RNA-seq analysis of LSK (Lin^-^Sca1^+^c-kit^+^) cells. **(A)** Principal component analysis in each genotype. **(B)** MA plot showing differentially expressed genes in each genotype (q < 0.05). Upregulated and downregulated genes are marked as red and blue circles, respectively. **(C)** Pathway analysis of double-MT-specific up-or down-regulated genes with CGP (chemical and genetic perturbations) gene sets in the Molecular Signatures Database (MSigDB). Enrichment is expressed as log_10_ FDR q-values. **(D)** Western blotting showing the levels of H2AK119Ub, H3K4me3, H3K27ac, H3K27me3, and total histone H3 in bone marrow LSK cells. **(E)** ChIP-seq analysis of bone marrow c-kit^+^ cells. The numbers of differential histone modifications at promoters (H3K27me3, H3K4me3, H3K27ac) and enhancers (H3K27ac) are shown (q < 0.05). **(F)** ATAC-seq analysis of bone marrow LSK cells. ATAC signals at promoters and enhancers (± 3kb) are shown. **(G)** HOMER motif analysis of differentially accessible promoters in each genotype. Representative HSC-related transcription factors highly ranked in double-MT mice are noted.

To investigate the mechanisms underlying the expression changes in double-MT mice, we assessed histone modifications in LSK cells by western blotting **(Figure 3D)**. The level of H2AK119Ub in Asxl1-MT KI and double-MT mice was decreased, which is consistent with a previous study that showed Asxl1-MT cooperates with Bap1 to deubiquitinate H2AK119Ub and de-repress gene expression ^29^. On the other hand, we did not observe an obvious difference in the levels of H3K4me3, H3K27me3, and H3K27ac among genotypes. To further evaluate histone modifications as well as chromatin accessibility in fine resolution, we performed chromatin immunoprecipitation sequencing (ChIP-seq) and assay for transposase-accessible chromatin sequencing (ATAC-seq) using bone marrow c-kit^+^ and LSK cells, respectively. The numbers of differential histone modifications (vs. control) for H3K27me3 and H3K4me3 at promoters were significantly increased in double-MT mice compared with Asxl1-MT KI or *Stag2* KO mice, whereas H3K27ac at promoters and enhancers was exclusively affected by *Stag2* loss **(Figure 3E)**. Chromatin accessibility at promoters and enhancers was moderately increased in Asxl1-MT KI mice, which was substantially enhanced by the concurrent *Stag2* loss (**Figure 3F**). Interestingly, binding motifs of HSC-related transcription factors, including *FLI1, PU.1, GABPA, ERG,* and *RUNX1*, were enriched in differentially accessible promoters in double-MT mice ^38–42^ (**Figure 3G**). These analyses revealed that concomitant mutations in *Asxl1* and *Stag2* profoundly alter histone modifications and chromatin accessibility, which would be responsible for perturbed gene expressions as well as HSC functions in double MT mice.

### Polycomb-regulated genes are susceptible to *Asxl1* and *Stag2* mutations

We next attempted to characterize genes that are subjected to *Asxl1* and *Stag2* mutations. To do so, all promoters were classified into 8 clusters (P1-P8) based on gene expression, chromatin accessibility, and binding intensities of histone modifications (H2AK119Ub, H3K27me3, H3K4me3, H3K27ac), Asxl1, and cohesin in normal bone marrow c-kit^+^ cells by means of unsupervised k-means clustering (**Figure 4A; left panel**). Each promoter was annotated with differential changes in gene expression, histone modifications, and chromatin accessibility (vs control) (**Figure 4A; right panel,** blue and red bars show downregulation and upregulation, respectively). The promoter clusters consist of inactive promoters (P1/P2) (**Figure 4A**; blue rectangle, low levels of gene expression, H3K4me3, and H3K27ac), active promoters (P3/P4/P5) (**Figure 4A**; red rectangle, high levels of gene expression, H3K4me3, H3K27ac), and polycomb-regulated promoters (P6/P7) (**Figure 4A**; green rectangle, high levels of H3K27me3 and H2AK119ub). P8 cluster was excluded from the subsequent analysis since there were only few promoters in this cluster. We found that P6 and P7 clusters (hereinafter referred to as “polycomb-regulated promoters”) were enriched with differentially expressed genes across all genotypes (**Figure 4B; green rectangle**). Of note, genes specifically upregulated in double-MT cells, but not in either Asxl1-MT or *Stag2* KO cells, were especially enriched in those with polycomb-regulated promoters (**Figure 4C**). Upregulation of these genes was accompanied by de-repression of promoters which is indicated by a decrease in H3K27me3 level and an increase in H3K4me3 and H3K27ac levels (**Figure 4D and 4E**). These findings indicate that polycomb-regulated genes are common targets of *Asxl1* and *Stag2* mutations, and that these mutations exert synergistic effects on their expressions.

**Figure 4.**
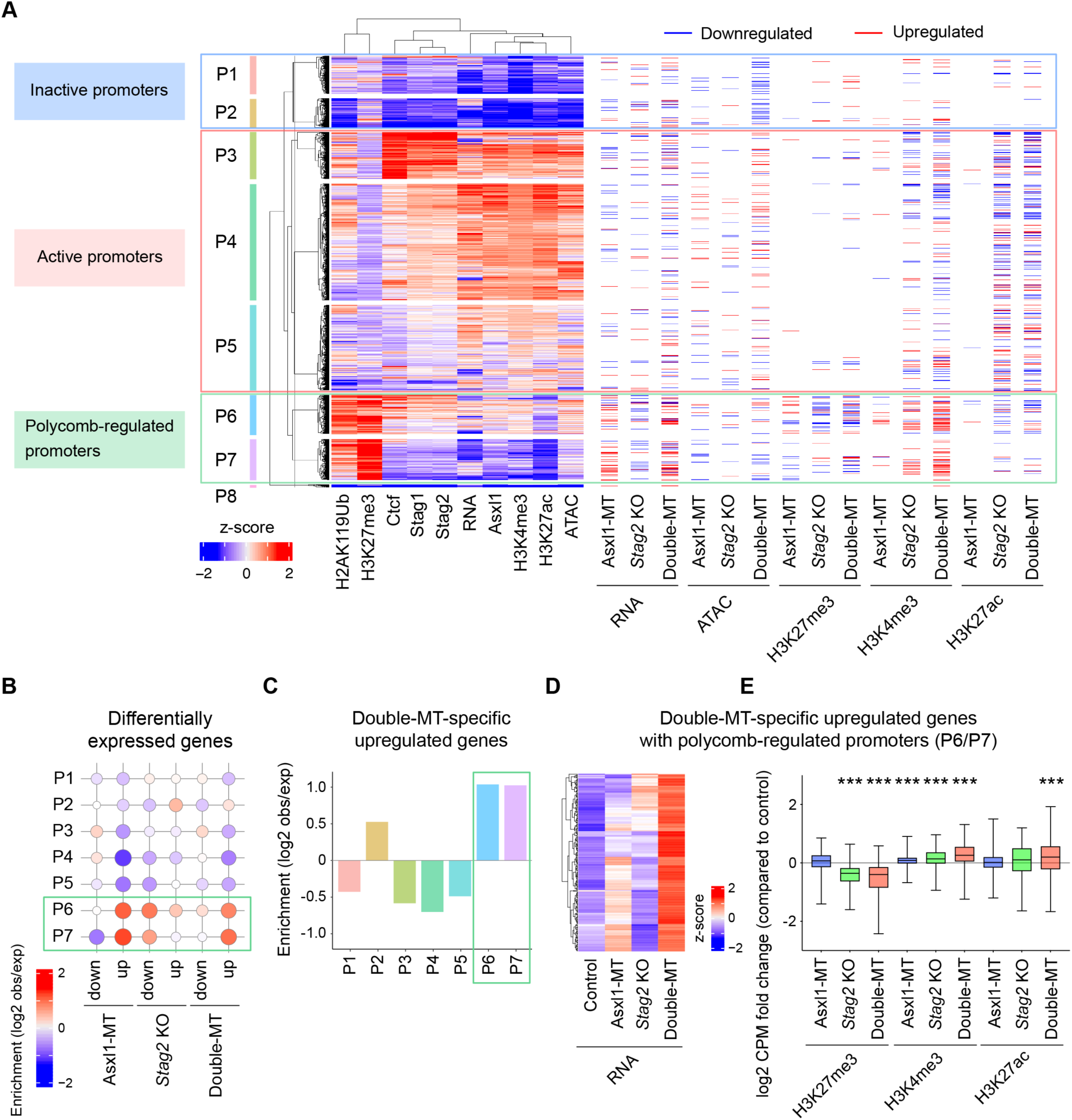
Polycomb-regulated promoters are susceptible to *Asxl1* and *Stag2* mutations. **(A)** Unsupervised k-means clustering of promoters in wild-type mice based on RNA-seq, ATAC-seq, and ChIP-seq (histone modifications, Asxl1, Stag1, Stag2, Ctcf) data. Signal intensities at each promoter are shown on the left panel. Differential expression, chromatin accessibility and histone modifications at each promoter are expressed as blue (downregulated) and red (upregulated) bars on the right panel. The promoter clusters are classified into inactive promoters (P1/P2) (blue rectangle, low levels of gene expression, H3K4me3, and H3K27ac), active promoters (P3/P4/P5) (red rectangle, high levels of gene expression, H3K4me3, H3K27ac), and polycomb-regulated promoters (P6/P7) (green rectangle, high levels of H3K27me3 and H2AK119ub). ChIP-seq datasets are publicly available at Sequence Read Archive (SRA) with the reference series tag “SRP133391” (Asxl1) and “SRP199092” (Stag1, Stag2, Ctcf). **(B)** Enrichment of differentially expressed genes in each promoter cluster. Differentially expressed genes are enriched in genes with polycomb-regulated promoters (P6/P7) (green rectangle). **(C)** Bar graph showing enrichment of double-MT-specific upregulated genes in each promoter cluster. Double-MT-specific upregulated genes are defined as genes whose expression levels are significantly upregulated in double-MT mice but not in Asxl1-MT KI or *Stag2* KO mice (see **Figure S4A**). Double-MT-specific genes are enriched in genes with polycomb-regulated promoters (P6/P7) (green rectangle). **(D)** Heatmaps indicating expression levels of double-MT-specific upregulated genes with polycomb-regulated promoters (P6/P7). **(E)** Fold changes of H3K27me3, H3K4me3, and H3K27ac signal levels at polycomb-regulated promoters (P6/P7) of double-MT-specific upregulated genes in each genotype (vs. control). Enrichment is defined as the ratio of observed (obs) number to expected (exp) number. Statistical significance was assessed by one-way ANOVA with Tukey–Kramer’s post-hoc test **(E)**. ***P < 0.001.

### Neither *Asxl1* nor *Stag2* mutations affect large-scale genome structure

In previous studies, we and others suggested that Stag2 plays a role in the maintenance of chromatin loops in hematopoietic cells, where it mediates short-range interactions within the genome, particularly P-E interactions ^23,35^. Thus, we performed Hi-C analysis of bone marrow c-kit^+^ cells to assess the effect of Asxl1-MT alone and in concert with *Stag2* KO on genome architecture (1.76 billion valid interactions per genotype on average). Globally, no large-scale alterations in the genome structure of compartmentalization (transcriptionally active A vs. inactive B compartments) or TADs were observed across all genotypes **(Figure S5A and S5B)**. Recapitulating the phenotype of *Stag2* KO cells as previously described, short-range interactions within individual TADs were substantially attenuated in *Stag2* KO and double-MT cells but hardly affected in Asxl1-MT KI cells **(Figure S5C)** ^23^.

### De-repression of polycomb-regulated genes induces ectopic interactions with active enhancers in the absence of *Stag2*

Given that genomic interactions between chromatin domains bearing specific histone marks occur upon cohesin removal ^12,13,21^, we hypothesized that ASXL1-MT-induced perturbation of histone modifications could affect chromatin architecture in the absence of Stag2. To test this possibility, we integrated RNA-seq, ChIP-seq, and Hi-C analyses to interrogate the impacts of histone modifications on P-E interactions and gene expression (**Figure 5A**). Furthermore, each promoter was annotated based on the aforementioned promoter classification to reveal the characteristics of promoters where P-E interactions are affected by *Asxl1* and *Stag2* mutations (**Figure 4A**).

**Figure 5.**
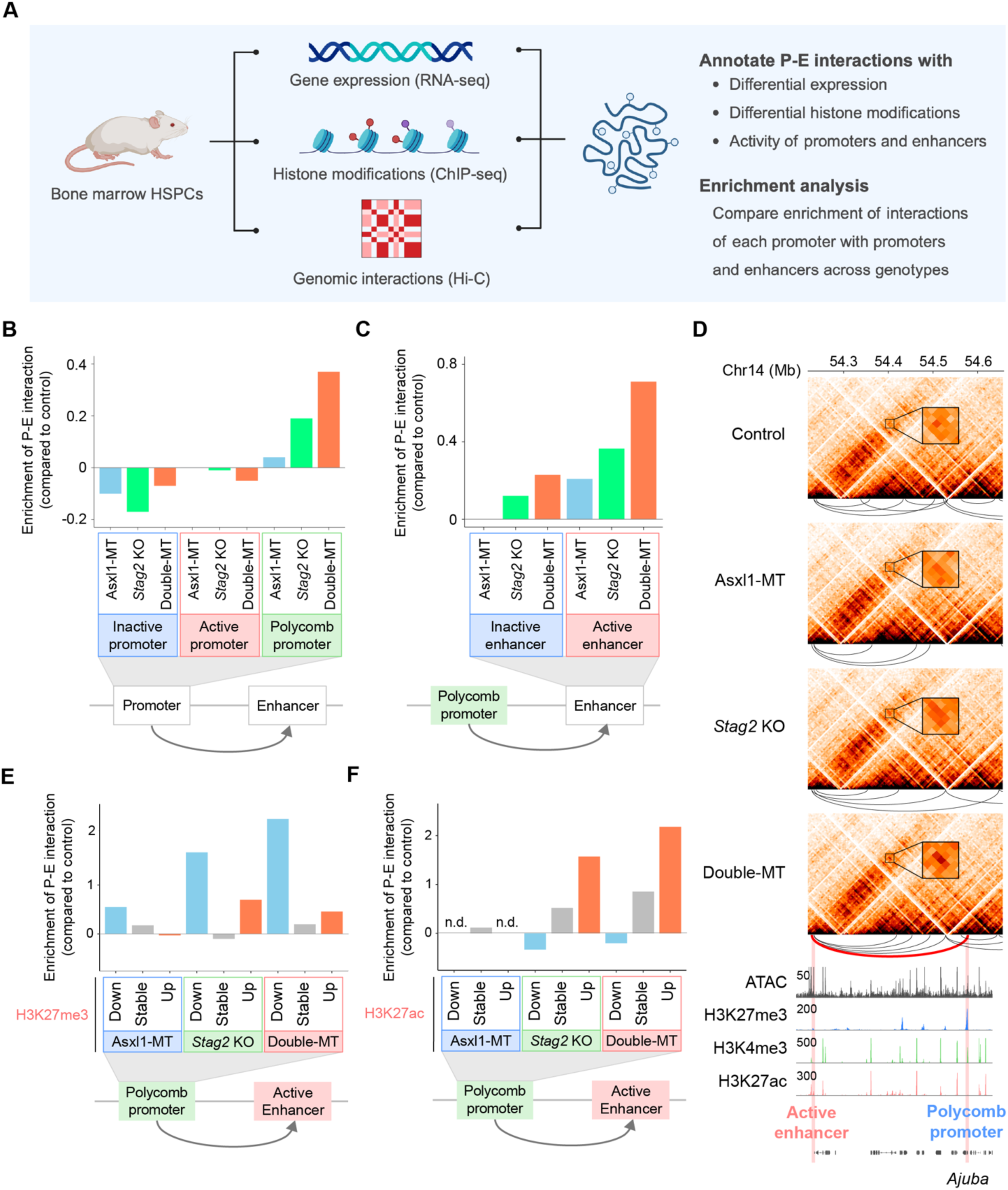
De-repression of polycomb-regulated promoters elicits genomic interactions with active enhancers in the absence of *Stag2*. (A) A scheme representing a workflow of multi-omics analysis. **(B)** Enrichment of P-E interactions at inactive, active, and polycomb-regulated promoters in each genotype (vs. control). **(C)** Enrichment of P-E interactions between polycomb-regulated promoters and inactive or active enhancers in each genotype (vs. control). **(D)** Representative contact map showing an ectopic interaction of polycomb-regulated promoter with active enhancer. An interaction between polycomb-regulated promoter of *Ajuba* gene and the active enhancer in double-MT cells is shown in a magnified image on contact map and a red heavy line below contact map. **(E, F)** Enrichment of P-E interactions between polycomb-regulated promoters with differential H3K27me3 **(E)** or H3K27ac **(F)** levels (vs. control) and active enhancers in each genotype (vs. control). n.d.; Data not available due to stable H3K27ac levels in Asxl1-MT cells. Enrichment is defined as the ratio of observed (obs) number to expected (exp) number. The difference in the enrichment of P-E interactions (vs. control) was calculated by subtracting the enrichment values of Asxl1-MT, *Stag2* KO, or double-MT cells from that of control cells. Negative and positive values represent depletion and enrichment compared with control mice, respectively.

This analysis demonstrated increased interactions between polycomb-regulated promoters and enhancers in double-MT cells and to a lesser extent in *Stag2* KO cells, but not in Asxl1-MT cells, indicating that polycomb-regulated promoters form Stag2-independent P-E interactions upon *Stag2* depletion **(Figure 5B)**. By contrast, P-E interactions at neither active promoters nor inactive promoters increased in double-MT cells, suggesting that the synergistic impacts of *Asxl1* and *Stag2* mutations on P-E interactions is specific for polycomb-regulated promoters. P-E interactions at polycomb-regulated promoters were more frequently generated with active enhancers (H3K4me1^+^/H3K27ac^+^) than with inactive enhancers (H3K27ac^-^) ^43,44^ **(Figure 5C, 5D, and S6A**; see double-MT specific P-E interaction indicated in a magnified image on contact map and a red heavy line below contact map in **Figure 5D)**. Because disruption of histone modifications was a notable feature in double-MT cells (**Figure 3E**), we evaluated the effect of perturbed histone modifications on P-E interactions. Notably, decreased H3K27me3 and increased H3K27ac levels at polycomb-regulated promoters were associated with increased interactions with active enhancers (**Figure 5E and 5F**), but not with inactive enhancers (**Figure S6B and S6C**), in *Stag2* KO and double-MT cells. On the other hand, the relationship between H3K4me3 levels and P-E interactions was unclear (**Figure S6D**). These results indicate that the de-repression of polycomb-regulated promoters, which was suggested by decreased H3K27me3 levels, induces Stag2-independent interactions with active enhancers upon *Stag2* loss.

We then explored the impact of altered P-E interactions on gene expressions in double-MT cells. Polycomb-regulated promoters of upregulated genes increased interactions with active enhancers but not with inactive enhancers (**Figure S6E**; orange bars), whereas those of downregulated genes decreased interactions with enhancers in double-MT cells (**Figure S6E**; blue bars), suggesting the correlation of P-E interactions with gene expressions (**Figure S6E**). However, we also observed increased interactions between polycomb-regulated promoters and enhancers in genes whose expression did not change (**Figure S6E; gray bars**), implying that the engagement of promoters with enhancers is insufficient for gene upregulation. Since double-MT-specific upregulated genes were especially enriched in polycomb-regulated genes **(Figure 4C**; green rectangle), we examined P-E interactions of these genes across genotypes. Polycomb-regulated promoters of double-MT-specific upregulated genes increased interactions with active enhancers in double-MT cells, but in neither Asxl1-MT nor *Stag2* KO cells **(Figure S6F)**. Of note, upregulation of these genes was accompanied by decreased H3K27me3 levels and increased H3K27ac levels at promoters (**Figure 4D and 4E**). Collectively, these data suggest that de-repression of polycomb-regulated promoters drives interactions with active enhancers upon Stag2 loss, resulting in aberrant gene upregulation.

### De-repression alters local genomic architecture of polycomb-regulated genes in the absence of Stag2-mediaged genomic interactions

Our Hi-C analysis revealed enriched interactions of active promoters with (i) active promoters and (ii) active enhancers at baseline level **(Figure 6A; orange circles)**, which would contribute to high expressions of the relevant genes. Based on these findings, it is assumed that active promoters assemble with other active promoters and enhancers to form hubs composed of genes with high transcriptional activity, a concept that aligns with multiple previous reports ^45–47,50,52,54^. Given that de-repressed polycomb-regulated promoters formed interactions with active enhancers in double-MT cells (**Figure 5B-E**), we hypothesized that de-repression converts these promoters from a repressive state to an active state (i.e., active promoter-like state), thereby inducing interactions with active promoters.

**Figure 6.**
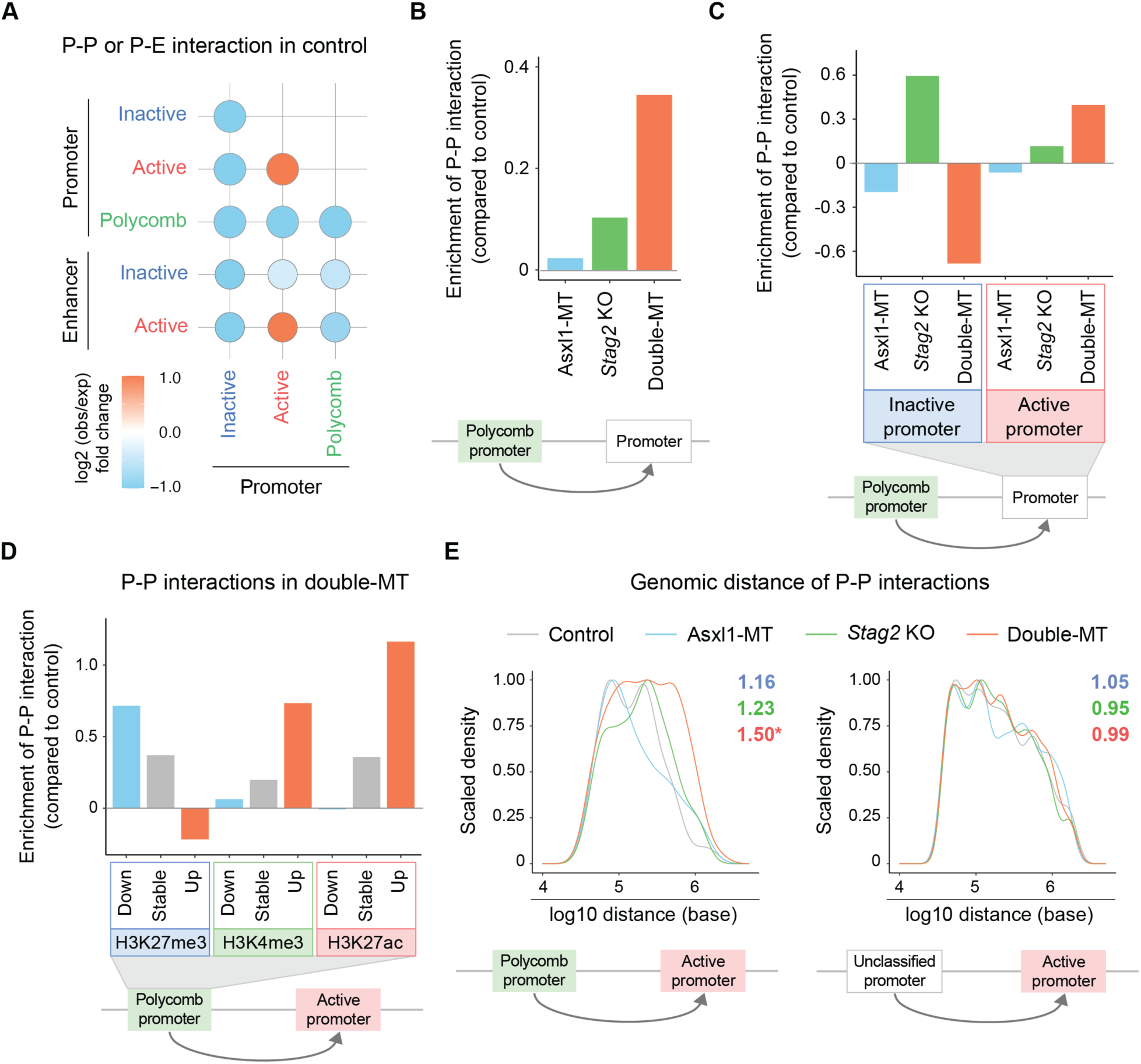
De-repression of polycomb-regulated promoters provokes engagement with active promoters accompanied by altered spatial arrangement of polycomb-regulated genes. **(A)** Enrichment of interactions between (i) inactive/active/polycomb-regulated promoters and (ii) inactive/active/polycomb-regulated promoters or inactive/active enhancers in control cells. **(B)** Enrichment of P-P interactions at polycomb-regulated promoters in each genotype (vs. control). **(C)** Enrichment of P-P interactions between polycomb-regulated promoters and inactive or active promoters in each genotype (vs. control). **(D)** Enrichment of P-P interactions at polycomb-regulated promoters with differential levels of H3K27me3, H3K4me3, or H3K27ac (vs. control) and active promoters in double-MT cells (vs. control). **(E)** Scaled density curve showing the genomic distances of P-P interactions between (i) polycomb-regulated promoters (left panel) or unclassified (all) promoters (right panel) and (ii) active promoters in each genotype. Fold changes in the average genomic distance of P-P interactions in ASXL1-MT (blue), *Stag2* KO (green), or double-MT (red) cells (vs. control) are noted. Statistical significances were assessed by pairwise comparisons using the Wilcoxon rank sum test with Benjamini-Hochberg correction. *P ≤ 0.05. Enrichment is defined as the ratio of observed (obs) number to expected (exp) number. Negative and positive enrichment values represent depletion and enrichment compared with the expected number of interactions, respectively. The difference in the enrichment of P-P interactions (vs. control) was calculated by subtracting the enrichment values of Asxl1-MT, *Stag2* KO, or double-MT cells from that of control cells. Negative and positive values represent depletion and enrichment compared with control mice, respectively.

Indeed, interactions of polycomb-regulated promoters with other promoters were increased in double-MT cells (**Figure 6B**), exclusively with active promoters (**Figure 6C**). In contrast, *Stag2* KO alone showed a lesser increase in interactions with promoters (**Figure 6B**), and this increase was restricted with inactive promoters (**Figure 6C**). These results suggest that Asxl1-MT-mediated de-repression of polycomb-regulated promoters is required for their engagement with active promoters. Consistent with this notion, de-repression of polycomb-regulated promoters, which was indicated by decreased levels of H3K27me3 and increased levels of H3K4me3 and H3K27ac, was associated with increased interactions with active promoters in double-MT cells **(Figure 6D)**. We next measured genomic distances of P-P interactions to explore the impacts of *Asxl1* and *Stag2* mutations on genomic conformation of polycomb-regulated genes. Polycomb-regulated promoters formed interactions with more distal active promoters on genome in double-MT cells compared with control, Asxl1-MT, and *Stag2* KO cells **(Figure 6E; left panel)**. This observation was unique to polycomb-regulated promoters since genomic distances of P-P interactions between unclassified promoters (all promoters) and active promoters did not change across genotypes (**Figure 6E; right panel**). These results suggest that de-repression of polycomb-regulated promoters altered local genomic structure independently of Stag2 cohesin-mediated genomic interactions, thereby forming transcription units composed of active promoters and active enhancers.

### Mutations in *ASXL1* and *STAG2* synergistically upregulate polycomb-regulated genes in human AML patients

Based on the above findings, we attempted to clarify the mechanistic link between our mouse model and human patients harboring mutations in *ASXL1* and *STAG2*. Taking advantage of the public transcriptome data of AML patients (Beat AML ^48^), we defined differentially expressed genes in patients with *ASXL1, STAG2,* or *ASXL1*/*STAG2* (hereinafter referred to as “double”) mutations compared to those with neither *ASXL1* nor *STAG2* mutations (hereinafter referred to as “AS wild-type”). We then quantified H3K27me3 levels at promoters in bone marrow CD34^+^ HSPCs from healthy donors ^48,49^ to identify polycomb-regulated genes **(Figure 7A)**. Expression changes in patients with double mutations (vs. AS wild-type) positively correlated with H3K27e3 levels at their promoters, which was more pronounced than those with *ASXL1* or *STAG2* single mutation **(Figure 7B)**. These findings suggest that polycomb-regulated genes are susceptible to concurrent mutations in *ASXL1* and *STAG2* in human patients.

**Figure 7.**
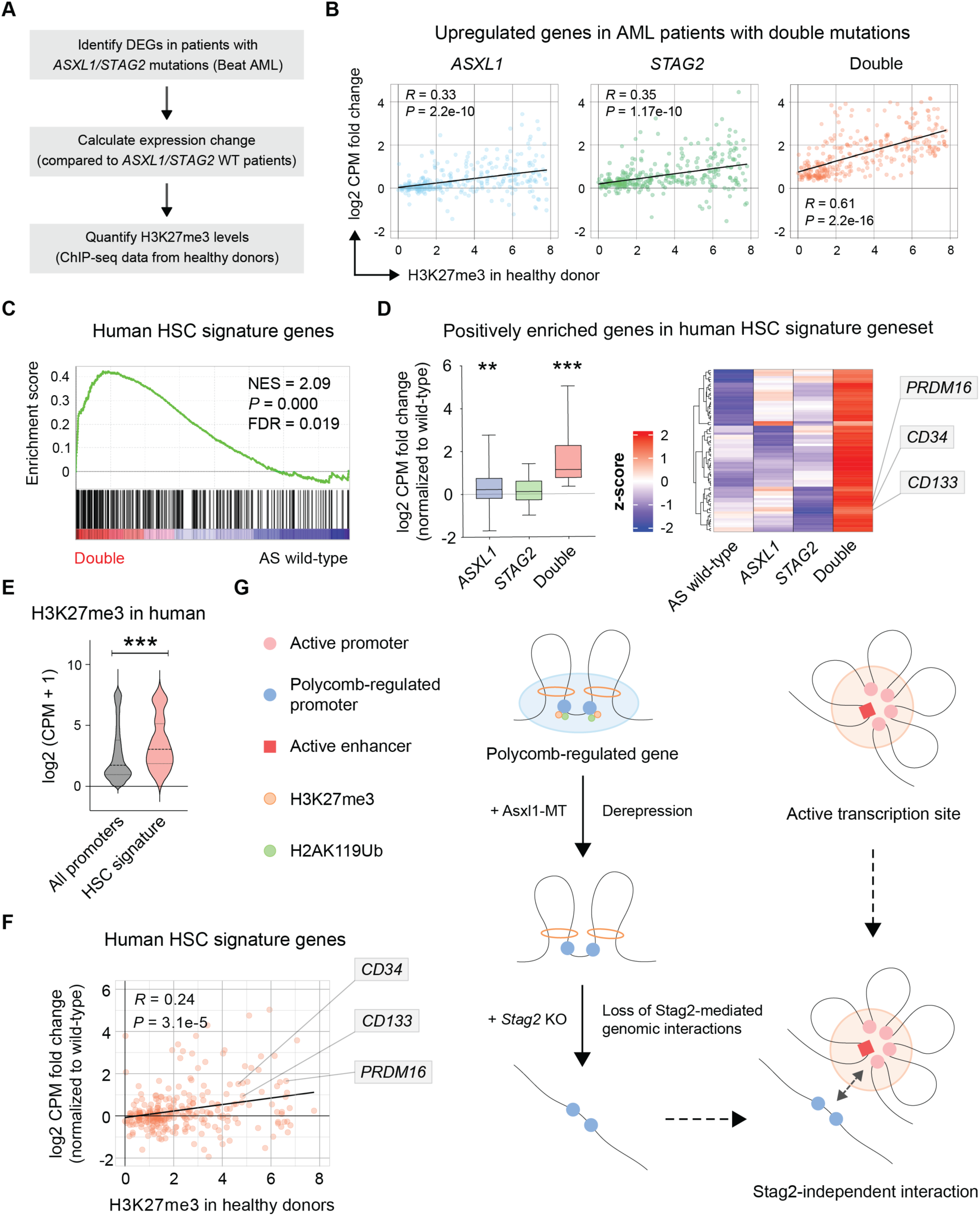
Mutations in *ASXL1* and *STAG2* cooperate to upregulate polycomb-regulated genes in patients with AML. **(A)** A flowchart of the analysis on transcriptomic data from human AML patients with *ASXL1* (N = 29), *STAG2* (N = 18), or *ASXL1/STAG2* (double) (N = 6) mutations and without either *ASXL1* or *STAG2* (N = 410) mutations. **(B)** Correlation of H3K27me3 levels (in healthy human) and expression changes of upregulated genes in AML patients with double mutations (vs. *ASXL1*/*STAG2* wild-type). *P* and *R* values were determined by a linear regression model. **(C)** GSEA indicating upregulation of HSC signature genes (Jaatinen et al., 2006) in AML patients harboring double mutations (vs. *ASXL1*/*STAG2* wild-type) **(D)** Expression changes of core enriched HSC signature genes in **(C)** (vs. *ASXL1*/*STAG2* wild-type). **(E)** H3K27me3 levels at promoters of human HSC signature genes and all promoters. **(F)** Correlation between H3K27me3 levels (in healthy human) and expression changes in core enriched HSC signature genes from **(C)** in AML patients with double mutations (vs. *ASXL1*/*STAG2* wild-type). *P* and *R* values were determined by a linear regression model. **(G)** A schematic diagram presenting cooperative effects of Asxl1-MT and *Stag2* loss on genomic interactions of polycomb-regulated genes. Asxl1-MT expression and *Stag2* depletion affect expression of polycomb-regulated genes through de-repression and disruption of genomic interactions, respectively. Asxl1-MT-mediated de-repression of polycomb-regulated promoters drives ectopic interactions with active enhancers upon *Stag2* loss. De-repressed polycomb-regulated promoters also organize interactions with active promoters that are regulated by active enhancers, resulting in their translocation from transcriptionally repressive sites to active sites.

We further sought to obtain the implication of the de-repression of polycomb-regulated genes for the pathogenesis of double mutation-driven myeloid malignancies by gene set enrichment analysis (GSEA). We found that HSC signature genes were significantly upregulated in double-MT mice and human AML patients with double mutations **(Figure 7C and S7A)**, an observation that would underlie the abnormal expansion of HSCs and development of MDS in double-MT mice (**Figure 2B-D and 2F-G**). Importantly, expression levels of HSC signature genes, including *PR domain containing 16 (PRDM16)*, *CD34*, and *CD133*, were much higher in patients with double mutations compared with those with *ASXL1* or *STAG2* mutation alone **(Figure 7D)**. Promoters of HSC signature genes commonly exhibited high H3K27me3 levels (vs. all promoters) in HSPCs from control mouse and healthy human (**Figure 7E and S7B**), suggesting that de-repression underlies upregulation of these genes in double-MT mice and human AML patients with double mutations. Indeed, Asxl1-MT expression and *Stag2* loss synergistically increased the expression levels of HSC signature genes accompanied by decreased level of H3K27me3 and increased levels of H3K4me3 and H3K27ac **(Figure S7C)**. Furthermore, expression changes of HSC signature genes in AML patients with double mutations were positively correlated with baseline H3K27me3 levels at promoters (**Figure 7F**), implying a possible link between de-repression of polybomb-regulated promoters and upregulation of genes involved in HSC functions. Taken together, these results suggest that *ASXL1* mutations corporates with *STAG2* mutations to upregulate polycomb-regulated genes, including HSC signature genes, resulting in myeloid transformation.

## DISCUSSION

Through a multi-omics approach, we here demonstrated that Asxl1-MT and *Stag2* loss synergistically perturb the gene expression, histone modifications, and genomic conformations of polycomb-regulated genes. Asxl1-MT expression and *Stag2* loss have been shown to dysregulate histone modifications and genome architecture, respectively, to perturb gene expression ^20,29–31,33,51^. Upon the combination of these two mutations, polycomb-regulated promoters were de-repressed and organized ectopic interactions with active enhancers, resulting in increased gene expression. These de-repressed polycomb-regulated promoters also organized interactions with distal active promoters which could be regulated by active enhancers. Therefore, we speculate that the de-repression of polycomb-regulated promoters augments their affinity to active promoters and active enhancers, translocating the polycomb-regulated genes from transcriptionally repressive sites to active sites in the absence of Stag2-mediated genome organization (**Figure 7G**).

We propose that the Stag2-mediated genome architecture constrains histone modification-driven, Stag2-independent genomic interactions, ensuring the robustness of gene expression. Previous studies have shown that chromosomal interactions at promoters are positively correlated with active histone marks and negatively correlated with repressive histone marks ^46,53^. Extending these findings, our study suggests that altered histone modifications at promoters actively change the combination of their interacting partners, an effect abrogated by the cohesin-mediated genome architecture. In the disease setting, the simultaneous disruption of histone modifications and the chromatin architecture can provoke Stag2-independent chromatin interactions to perturb gene expressions, manifesting the significance of concomitant mutations in the epigenetic regulator and Stag2-cohesin complex in oncogenesis.

## METHODS

### Mice

Wild-type C57BL/6J mice were bred in-house. Conditional Asxl1-MT KI and *Stag2* KO mice were generated as previously described and crossbred with *Vav-Cre* transgenic mice ^23,31^. All experiments were performed with 6-12-week-old mice. The mice were housed using a 12-hour dark/light cycle at 20-25 °C and a humidity between 40% and 60%. The experiments were approved by the Committee on the Ethics of Animal Experiments, and all mice were maintained according to the guidelines of the Institute of Laboratory Animal Science (PA13–19 and PA16–31).

### Flow cytometry

Bone marrow cells were isolated by flushing long bones (femurs and tibias) in phosphate-buffered saline (PBS) containing 2% heat-inactivated FBS (FACS buffer). Cell suspensions were lysed with erythrocyte lysis buffer (150 mM NH_4_Cl, 10 mM KHCO_3_, 100 μM EDTA-Na_2_), filtered through a 40 μm filter, and incubated with a cocktail of biotinylated monoclonal antibodies to lineage markers (CD5, B220, CD11b, Gr-1, and Ter119) and then anti-biotin microbeads (Miltenyi Biotec). Lin^−^ cells were purified using MACS separation LS columns (Miltenyi Biotec). To stain HSCs, cells were then stained with CD150-PE (BioLegend, 115904), c-kit-PE-Cy7 (BioLegend, 105814), Sca1-APC (BioLegend, 108112), CD48-Brilliant Violet 421 (BioLegend, 103427), and Streptavidin-Brilliant Violet 605 (BioLegend, 405229). To stain myeloid progenitors, CD16/32-PE (BD, 553145) and CD34-eFluor450 (Thermo Fisher Scientific, 48-0341-82) were used instead of CD150-PE and CD48-Brilliant Violet 421. For the apoptosis and cell cycle analysis, CD48-APC-Cy7 (BioLegend, 103431) and Sca1-Brilliant Violet 785 (BioLegend, 108139) antibodies were used instead of CD48-Brilliant Violet 421 and Sca1-APC antibodies. To assess erythroid differentiation, cells harvested from bone marrow and spleen were stained with CD71-PE (BD, 553267) and Ter119-PE/Cy7 (BioLegend, 116222). To stain differentiation markers, peripheral blood cells and bone marrow cells were analyzed using CD11b-PE (eBiosciense, 12-0112-85), CD3-APC (BioLegend, 100236), and CD45R/B220-APC/Cy7 (BioLegend, 103224) antibodies. Propidium iodide (PI) or 4′,6-diamidino-2-phenylindole (DAPI) was used to exclude dead cells from the analysis. All data were collected using FACSVerse or FACSAria.

### Western blotting

FACS-sorted LSK cells were denatured with Laemmli sample buffer (Bio-Rad Laboratories) at 95C° for 5 min. Lysates of 30,000 LSK cells were subjected to electrophoresis, transferred to nitrocellulose membranes, and blocked with 5% bovine serum albumin (BSA)/TBS for 1 hour at room temperature. Samples were then incubated with primary antibodies in 1% BSA/0.1% Tween 20/TBS overnight at 4°C, washed and incubated with secondary antibodies in 1% BSA/0.1% Tween 20/TBS for 1 hour at room temperature. The primary antibodies used in the immunoblotting were anti-H3K4me3 (clone CMA304, 1;1000 (kindly provided by Dr. Hiroshi Kimura)), anti-H3K27me3 (clone CMA323, 1:1000 (kindly provided by Dr. Hiroshi Kimura)), anti-H2AK119Ub (Cell Signaling Technologies, #8240, 1:1000), and anti-H3K27ac (Cell Signaling Technologies, #8173, 1:500).

### Serial transplantation

1.0 × 10^6^ bone marrow cells harvested from Asxl1-MT KI, *Stag2* KO, or double-MT mice (CD45.2^+^) were mixed with 1.0 × 10^6^ wild-type bone marrow cells and intravenously injected into lethally irradiated (900 cGy total, divided into two doses of 450 cGy given over 4 h) wild-type recipient mice (CD45.1^+^). Six months after the transplantation, 6 × 10^6^ million bone marrow cells were isolated from three recipient mice and serially transplanted into lethally irradiated recipients.

### Cell cycle analysis

Bone marrow cells stained with antibodies for surface markers were fixed in 4% paraformaldehyde for 10 min at room temperature, permeabilized with 0.2% Triton X-100 for 15 min at room temperature and blocked with 5% goat serum for 15 min at room temperature. The cells were then stained with Ki-67-eFluor660 (eBioscience, 50-5698-82) and DAPI for 30 min at 4 °C.

### Apoptosis analysis

Bone marrow cells were stained with surface marker antibodies and then with Annexin V-APC (BioLegend, 640920) antibody in binding buffer (10 mM HEPES (pH 7.4), 140 mM NaCl, 2.5 mM CaCl2) for 30 min at room temperature.

### Mutation analysis in human MDS/AML

We analyzed the combination of mutations in human MDS/AML patients using in-house or publicly available data of large-scale genetic profiling of MDS/AML ^23,55–61^. Patient survival and clinical characteristics were further investigated in 831 MDS patients ^57^. Pairwise comparisons were performed by the Wilcoxon test for continuous variables. Overall survival was estimated using the Kaplan–Meier method and compared among groups with the log-rank test. Significance was determined using a two-sided α level of 0.05. Multiple testing was adjusted according to the method described by Benjamini and Hochberg ^62^.

### RNA-sequencing

RNA-seq experiments were performed in three biological replicates. Total RNA was isolated from sorted LSK cells using an RNeasy Micro Kit (QIAGEN). RNA samples with RIN > 9 were proceeded to the sequencing analysis. The synthesis and amplification of cDNA was performed using an NEBNext Ultra RNA Library Prep Kit for Illumina (New England Biolabs), followed by high-throughput sequencing on a HiSeq 2500 (Illumina) with 126 bp paired-end reads. The sequencing reads were aligned to the reference genome (mm9) using STAR (v2.5.3). Reads on each refSeq gene were counted with the script annotatePeaks.pl from the HOMER package ^63^, and the edgeR package was used to identify differentially expressed genes with a q-value threshold of 0.05 ^64^. The analysis was performed in genes expressed at > 1 count per million (CPM) in two or more samples, and generalized linear models were used to compare gene expression data. A gene set enrichment analysis (GSEA) was performed with the GSEA tool from the Broad Institute (http://software.broadinstitute.org/gsea/). To classify differentially expressed genes in double-MT mice, double-MT-specific down/upregulated genes were defined as genes whose expression levels are down/upregulated in double-MT mice but not in either Asxl1-MT KI or *Stag2* KO mice. Asxl1-MT-dependent down/upregulated genes were defined as genes whose expression levels are down/upregulated in Asxl1-MT KI and double-MT mice but not in *Stag2* KO mice. *Stag2* KO-dependent down/upregulated genes were defined as genes whose expression levels are down/upregulated in *Stag2* KO and double-MT mice but not in Asxl1-MT KI mice.

### ATAC-sequencing

ATAC-seq experiments were performed in biological duplicates as previously described ^23^. Briefly, freshly isolated 10,000 LSK cells were pelleted, and 50 μL transposase mixture (25 μL of 2 x TD buffer, 2.5 μL TDE1, 0.5 μL of 1% digitonin, and 22 μL nuclease-free water) (FC-121-1030, Illumina; G9441) was added to the cells. After transposition reactions at 37°C for 30 min, transposed DNA was purified using a QIAGEN MinElute Reaction Cleanup Kit. Transposed fragments were PCR-amplified, and the resulting library was sequenced on a NoveSeq 6000 (Illumina). Reads were aligned to the mouse mm9 reference genome using bowtie2 (v2.3.3) with-X 2000 –no-mixed –very-sensitive parameters following adapter trimming using cutadapt (v1.14). Duplicates were removed using Picard (v2.6.0), and reads on mitochondria genome or blacklisted regions (ENCODE) were removed using bedtools (v2.27.1). Peaks were called with MACS (v2.1.1) using –nomodel –broad parameters and a q-value threshold of 1 x 10^-5^ for individual replicates as well as the merged data of all samples ^65^. Reads on each promoter and enhancer were counted with the script annotatePeaks.pl from the HOMER package, and the edgeR package was used to identify the differentially accessible promoters and enhancers with a p-value threshold of 0.05 ^64^. A motif analysis of differentially accessible enhancers was conducted using the script findMotifsGenome.pl from the HOMER package ^63^.

### ChIP-sequencing

ChIP-seq experiments were performed in biological duplicates as previously described ^23^. Freshly isolated c-kit^+^ cells were fixed in PBS with 1% formaldehyde (Thermo Fisher Scientific) for 10 min at room temperature with gentle mixing. The reaction was stopped by adding glycine solution (10x) (Cell Signaling Technology) and incubating for 5 minutes at room temperature, and the cells were washed in cold PBS twice. The cells were then processed with a SimpleChIP Plus Sonication Chromatin IP Kit (Cell Signaling Technology) and Covaris E220 (Covaris) according to the manufacturer’s protocol. The antibodies used for ChIP are as follows: H3K27ac (Cell Signaling Technology, D5E4), H3K27me3 (Cell Signaling Technology, C36B11), H2AK119Ub (Cell Signaling Technology, D27C4), or H3K4me3 (Cell Signaling Technology, C42D8). After purification of ChIPed DNA, ChIP-seq libraries were constructed using a ThruPLEX DNA-seq Kit (Takara) according to the manufacturer’s protocol and then subjected to sequencing using a NoveSeq 6000. ChIP-seq experiments were performed using two biological replicates with input controls. The sequencing reads were aligned to the reference genome (mm9) using bowtie (v1.2.2) following trimming of adapters and read tails to a total length of 50 base pairs using cutadapt. Duplicates and reads on blacklisted regions (ENCODE) were removed by Picard and bedtools, respectively. Reads on each promoter and enhancer were counted with the script annotatePeaks.pl from the HOMER package ^63^, and the edgeR package was used to identify differential ChIP-seq signals at promoters and enhancers with a q-value threshold of 0.05^64^. We also used ChIP-seq datasets available at Sequence Read Archive (SRA) with the reference series tag “SRP133391” (Asxl1), “SRP199092” (Stag1, Stag2, Ctcf), and “SRP060558” (human H3K27me3).

### k-means clustering

All promoters (≤ 2 kb from TSS) and enhancers (≤ 1 kb from ATAC peaks excluding ≤ 2 kb from TSS) were classified by k-means clustering according to the signal intensities of RNA-seq, ChIP-seq (histone modifications (H2AK119Ub, H3K27me3, H3K4me3, and H3K27ac for promoters, and H3K4me1 and H327ac for enhancers), Asxl1, Stag1, Stag2, and Ctcf), and ATAC-seq with the “ComplexHeatmap” package in R software^66^. The optimized number of clusters was determined using the “clusGap” package in R software ^67^. Promoters and enhancers with specific genomic features (differential expression, accessibility, and histone modifications) were assigned to the corresponding clusters, and their enrichment for each cluster was expressed as follows:

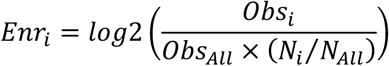

*Enr_i_* is the enrichment of genomic features in cluster *i*, *Obs_i_* is the observed genomic features in cluster *i*, *Obs_All_* is the number of all genomic features at promoters or enhancers, *N_i_* is the number of promoters or enhancers in cluster *i*, and *N_All_* is the number of all promoters or enhancers. Negative and positive values represent depletion and enrichment compared with the expected number of genomic features, respectively.

### Hi-C

Hi-C experiments were performed using MboI restriction enzyme as previously described ^6^. Briefly, two million mouse c-Kit^+^ HSPCs were crosslinked with 1% formaldehyde for 10 min at room temperature. Cells were permeabilized and chromatin was digested with MboI restriction enzyme, and the ends of the restriction fragments were labeled with biotinylated nucleotides and ligated. After crosslink reversal, DNA was purified and sheared with Covaris M220 (Covaris). Then, point ligation junctions were pulled down with streptavidin beads. Libraries were constructed using the Nextera Mate Pair Sample Preparation Kit (Illumina) according to the manufacturer’s protocol and subject to sequencing using a NovaSeq 6000. Hi-C experiments were performed in biological duplicates. The sequencing reads were processed using Juicer ^6^ and mm9 reference genome ^6^. For a comparative analysis, valid interactions for each sample after filtering were set to equal the lowest number of interactions detected by randomly resampling the reads. The resulting average valid interactions per genotype resulted in 1.76 billion. Contact matrices used for further analysis were created for each replicate as well as for merged replicates by genotype and Knight-Ruiz (KR)-normalized with Juicer. Genomic compartmentalization (A or B compartments) was analyzed using eigenvectors at 25 kb resolution ^6^. A compartments were assigned to the genomic bin with positive eigenvector values and higher gene density, and B compartments were assigned to the genomic bin with negative eigenvector values and lower gene density. Loops were called at 5 kb resolution using Mustache with a p-value threshold of 0.05 ^68^ and then merged to construct loop sets. We performed a hierarchical TAD analysis using rGMAP at 5 kb resolution with dom_order = 3 ^69^ to identify hierarchical TAD structures such as TADs (level 1) and sub-TADs (level 2/3). An aggregated TAD analysis was performed separately according to the TAD levels. We resized each TAD into a 100 x 100 submatrix and calculated the sum of the size-normalized submatrices. Hi-C contact matrices were visualized using HiCExplorer ^70^.

### Analysis of genomic interactions

To investigate the effects of Asxl1-MT and *Stag2* loss on the chromatin architecture, anchors of genomic interactions were annotated with promoters (≤ 2 kb from TSS) and enhancers (≤ 1 kb from ATAC peaks excluding ≤ 2 kb from TSS) to identify promoter-enhancer (P-E) and promoter-promoter (P-P) interactions using the GenomicInteractions package in R/Bioconductor^71^. The anchors belonging to promoters and enhancers were assigned to the clusters defined by k-means clustering and enumerated in each cluster. We computed the enrichment of all pairwise inter-cluster interactions in each genotype under the assumption of a uniform distribution of P-E (or P-P) interactions across all promoters and enhancers. Enrichment is expressed as follows:

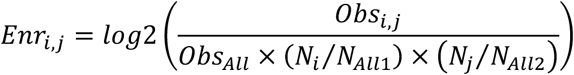

*Enr_i,j_* is the enrichment of the interactions between promoter cluster *i* and enhancer (or promoter) cluster *j*, *Obs_i,j_* is the observed interactions between promoter cluster *i* and enhancer (or promoter) cluster *j*, *Obs_All_* is the number of all P-E (or P-P) interactions, *N_i_* is the number of promoters in cluster *i*, *N_All1_* is the number of all promoters, *N_j_* is the number of enhancers (or promoters) in cluster *j*, and *N_All2_* is the number of all enhancers (or promoters). Negative and positive values represent depletion and enrichment compared with the expected number of interactions, respectively. The difference in the enrichment of P-E and P-P interactions was calculated by subtracting the enrichment values of Asxl1-MT KI, *Stag2* KO, or double-MT mice from that of control mice. Negative and positive values represent depletion and enrichment compared with control mice, respectively.

### Assessment of correlation between genomic features and interactions

To explore the relationship between genomic profiles and chromosomal architectures, anchors annotated with promoters and enhancers were further linked to genomic features (differential expression, differential histone modifications, binding status of Asxl1 and Stag2). We then calculated the enrichment of P-E (or P-P) interactions at promoters or enhancers with specific features under the assumption where all P-E (or P-P) interactions are uniformly distributed across all promoters and enhancers. The enrichment of interactions between promoters with feature A and enhancers (or promoters) with feature B is expressed as follows.

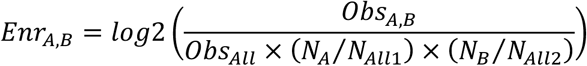

*Enr_A,B_* is the enrichment of interactions between promoters with feature A and enhancers (or promoters) with feature B, *Obs_A,B_* is the observed interactions between promoters with feature A and enhancers (or promoters) with feature B, *Obs_All_* is the number of all P-E (or P-P) interactions, *N_A_* is the number of promoters with feature A, *N_All1_* is the number of all promoters, *N_B_* is the number of enhancers (or promoters) with feature B, and *N_All2_* is the number of all enhancers (or promoters). Then, we calculated the difference in the enrichment of P-E or P-E interactions by subtracting the enrichment values of Asxl1-MT KI, *Stag2* KO, or double-MT mice from that of control mice, assessing the correlation of genome interactions with the binding status of Asxl1 and/or Stag2, expression changes, and differential histone modifications.

### Analysis of transcriptome data of human AML patients

RNA-seq data of human AML patients was obtained from a previous report ^48^. The sequencing reads were aligned to the reference genome (hg19) using STAR (v2.5.3). Reads on each refSeq gene were counted with featureCounts (v1.5.3) from the Subread package, and the edgeR package in R was used to identify differentially expressed genes with an FDR threshold of 0.05 ^64^. The analysis was performed in genes expressed at > 1 count per million (CPM) in two or more samples, and generalized linear models were used to compare gene expression data.

## ACKNOWLEDGMENTS

This work was supported by Japan Society for the Promotion of Science (Grant-in-Aid for Scientific Research (B) (No. 23K27442) (to H.H.), Grant-in-Aid for Scientific Research on Innovative Areas (No. 19H05745) (to H.K.), Grant-in-Aid for Transformative Research Areas (A) (No. 20H05940) (to K.S.), Grant-in-Aid for Scientific Research (S) (No. 20H05686) (to K.S.), Grant-in-Aid for Scientific Research (B) (No. 23K24360) (to S.G.), Grant-in-Aid for Scientific Research on Innovative Areas (No. 15H05909) (to S.O.), Grant-in-Aid for Scientific Research (S) (No. 19H05656) (to S.O.), Grant-in-Aid for Specially Promoted Research (No. 24H00009) (to S.O.), Grant-in-Aid for Scientific Research (A) (No. 20H00537) (to T.K.), Grant-in-Aid for Research Activity Start-up (No. 20K22809) (to Y.O.), Grant-in-Aid for Early-Career Scientists (No. 22K16320, 24K19223) (to Y.O.)), Japan Agency for Medical Research and Development (AMED) ((No. JP24gm1310006 (to H.H.), JP21cm0106501h0006 (to S.O.), JP19ck0106250h0003 (to S.O.)), Moonshot Research and Development Program (No. JP22zf0127008 (to H.K.), JP22zf0127009 (to S.O.)), the Ministry of Education, Culture, Sports, Science and Technology of Japan (the High Performance Computing Infrastructure System Research Project (No. hp160219, hp170227, hp180198, hp190158, hp200138, hp210167, JPMXP1020200102) (to S.O. and S.M.)), the Japanese Society of Hematology Research Grant (to T.K.), the Japan Science and Technology Agency (JST) (Fusion Oriented REsearch for disruptive Science and Technology (FOREST) Program (No. JPMJFR220L) (to Y.O.)), Takeda Science Foundation (to S.O. and Y.O.), Daiichi Sankyo Foundation of Life Science (to Y.O.), Princess Takamatsu Cancer Research Fund (to Y.O.), Kanae Foundation for the Promotion of Medical Science (to Y.O.), Ichiro Kanehara Foundation for the promotion of Medical Sciences and Medical Care (to Y.O.), Mochida Memorial Foundation for Medical and Pharmaceutical Research (to Y.O.), Japan Leukemia Research Fund (to Y.O.). We thank S. Yabuta at Kyoto University for technical assistance and IMSUT FACS Core Laboratory for flow cytometry.

## AUTHOR CONTRIBUTIONS

Experiments were designed and data were interpreted by T.F. under the guidance of S.G., S.O., and T.K. All experiments were carried out by T.F. with assistance from Y.S. and S.S. except the analysis of MDS and AML patient data for Figure 1A and 1B, Figure 7, and Figure S1, which was done by Y.O. and T.M.; RNA-seq, which was done by A.K.; ChIP-seq and ATAC-seq, which were done by Y.O.; and the Hi-C analysis, which was done by Y.O., M.K., T.S., K.S., and S.M. H.H. generated ASXL1-MT KI mouse. H.K. and M.N. generated *Stag2* KO mouse. The manuscript was written by T.F. All authors discussed the results and commented on the manuscript.

---

**Figure S1.**
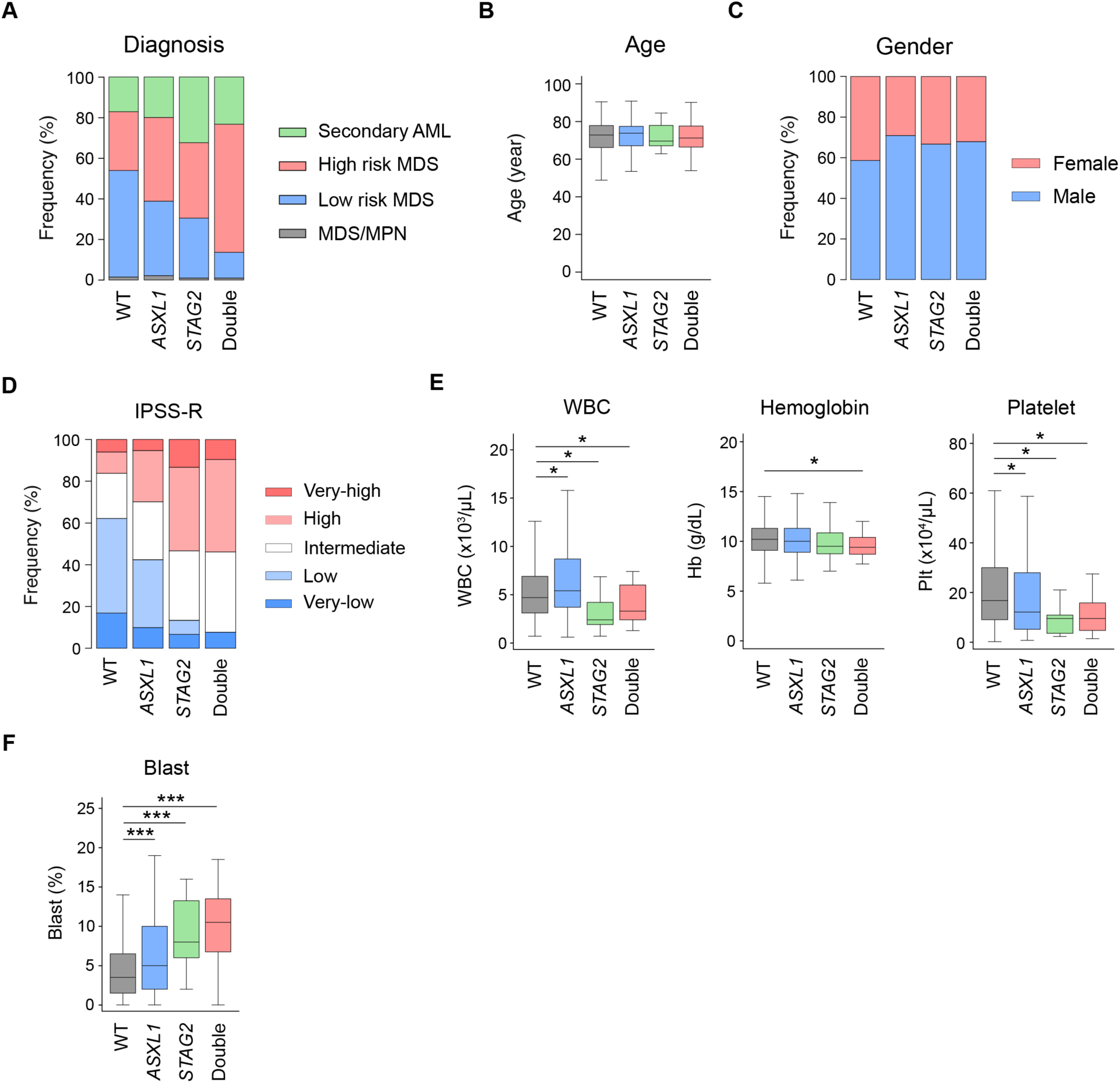
Characterization of MDS patients with mutations in *ASXL1* and *STAG2*. Clinical manifestation of MDS patients in our cohort (N = 831). **(A-D)** Diagnosis **(A)**, age **(B)**, gender **(C)**, and the Revised International Prognostic Scoring System (IPSS-R) **(D)** of the patients. For **(A)**, high-risk MDS includes RAEB-1 and RAEB-2. Other disease types are classified into low-risk MDS group. **(E)** Enumeration of white blood cells (WBCs), hemoglobin (Hb), and platelets (Plt). **(F)** The frequency of blast cells in WBCs. Statistical significances were assessed by one-way ANOVA with Tukey–Kramer’s post-hoc test. *P < 0.05, ***P < 0.001.

**Figure S2.**
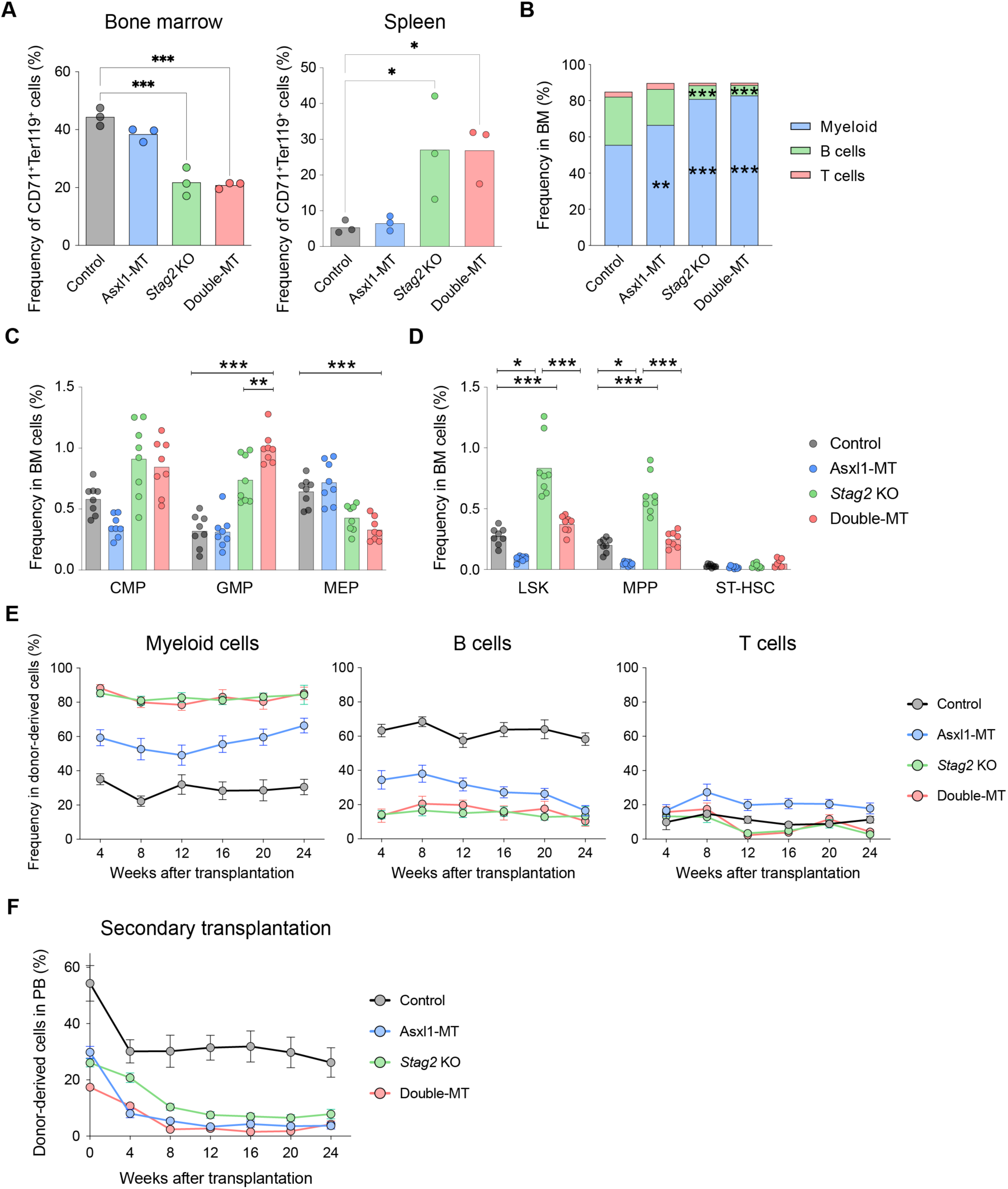
Asxl1-MT cooperates with *Stag2* loss to impair hematopoiesis. **(A)** The frequencies of erythroid progenitors (CD71^+^Ter119^+^) in bone marrow (left panel) and spleen (right panel) Bar graphs indicate mean value. **(B)** The frequencies of myeloid (CD11b^+^), B (B220^+^), and T (CD3^+^) cells in bone marrow. Bar graphs indicate mean value. **(C)** The frequencies of common myeloid progenitors (CMPs), granulocyte/macrophage progenitors (GMPs), megakaryocyte/erythroid progenitors (MEPs) in bone marrow. Bar graphs indicate mean value. **(D)** The frequencies of Lin^-^Sca1^+^c-kit^+^ (LSK) cells, multipotent progenitors (MPPs), and short-term hematopoietic stem cells (ST-HSCs) in bone marrow. Bar graphs indicate mean value. **(E)** The frequencies of myeloid (CD11b^+^), B (B220^+^), and T (CD3^+^) cells in donor-derived peripheral blood WBCs at the indicated weeks after primary transplantation. Error bars indicate mean values ± S.E. N = 12 (Control), 15 (Asxl1-MT), 12 (*Stag2* KO), or 15 (double-MT) mice/group. **(F)** 24 weeks after primary transplantation, secondary transplantation was performed with pooled 6 × 10^6^ bone marrow mononuclear cells isolated from three recipient mice. Donor chimerism in peripheral blood was assessed at the indicated weeks after secondary transplantation. Error bars indicate mean values ± S.E. N = 14 (Control), 11 (Asxl1-MT), 10 (*Stag2* KO), or 8 (double-MT) mice/group. Statistical significance was assessed by one-way ANOVA with Tukey–Kramer’s post-hoc test. *P < 0.05, **P < 0.01, ***P < 0.001.

**Figure S3.**
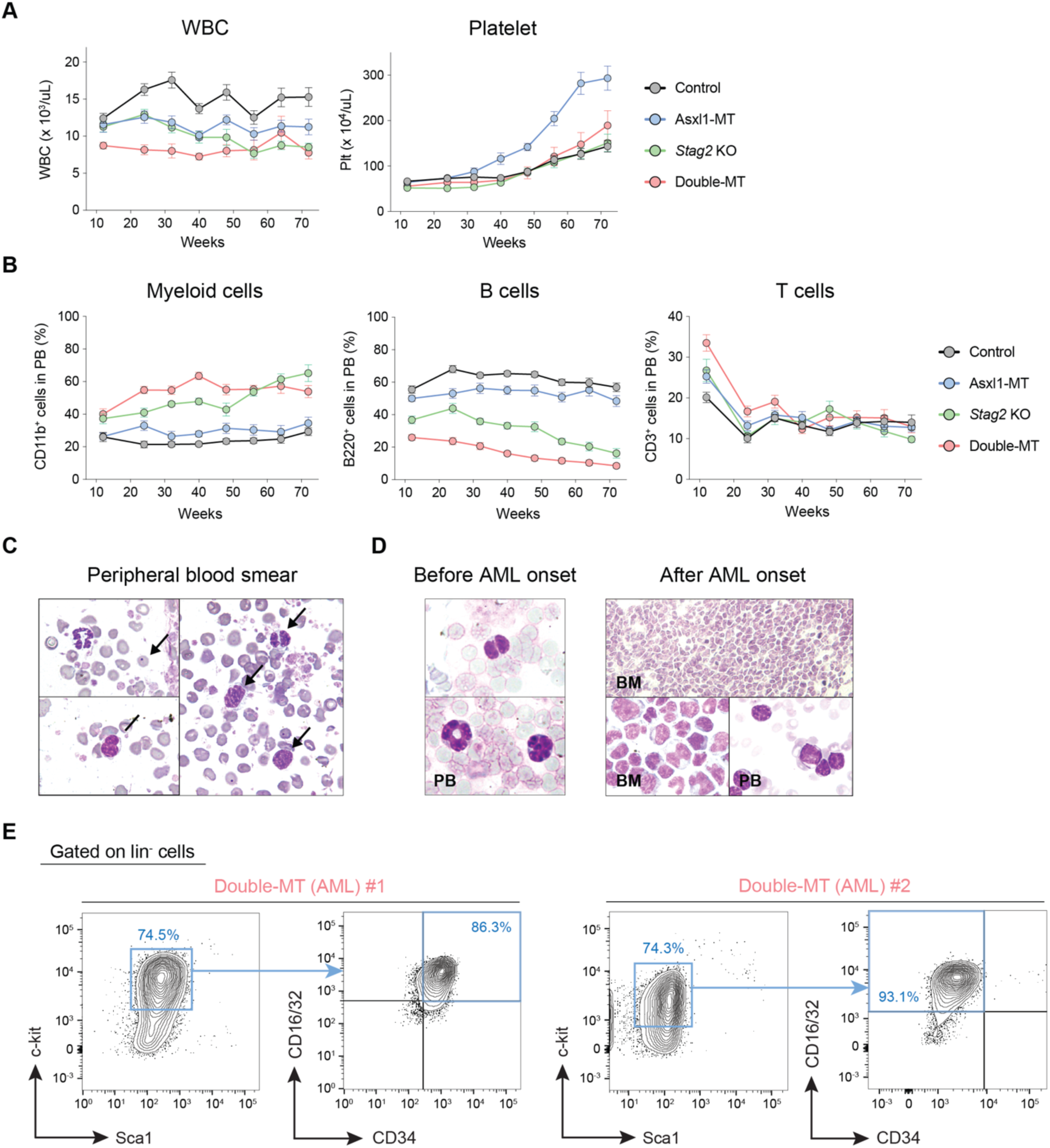
Double-MT mice developed MDS with hypocellular bone marrow. **(A)** Enumeration of white blood cells (WBCs) and platelets (Plt) at the indicated weeks after birth. N = 24 (Control), 21 (Asxl1-MT), 20 (*Stag2* KO), or 25 (double-MT) mice/group. Error bars indicate mean values ± S.E. **(B)** The frequencies of myeloid (CD11b^+^), B (B220^+^), and T (CD3^+^) cells in peripheral blood (PB) at the indicated weeks after birth. N = 24 (Control), 21 (Asxl1-MT), 20 (*Stag2* KO), or 25 (double-MT) mice/group. Error bars indicate mean values ± S.E. **(C)** Wright-Giemsa staining of peripheral blood smear from double-MT mice at the preclinical stage of MDS. **(D)** Wright–Giemsa staining of peripheral blood smear and bone marrow cells in double-MT mice before (left panel) and after (right panel) AML onset. **(E)** Representative plots of bone marrow HSPCs showing the expansion of GMPs (left panel) and L-GMP-like populations (right panel) in double-MT mice with AML.

**Figure S4.**
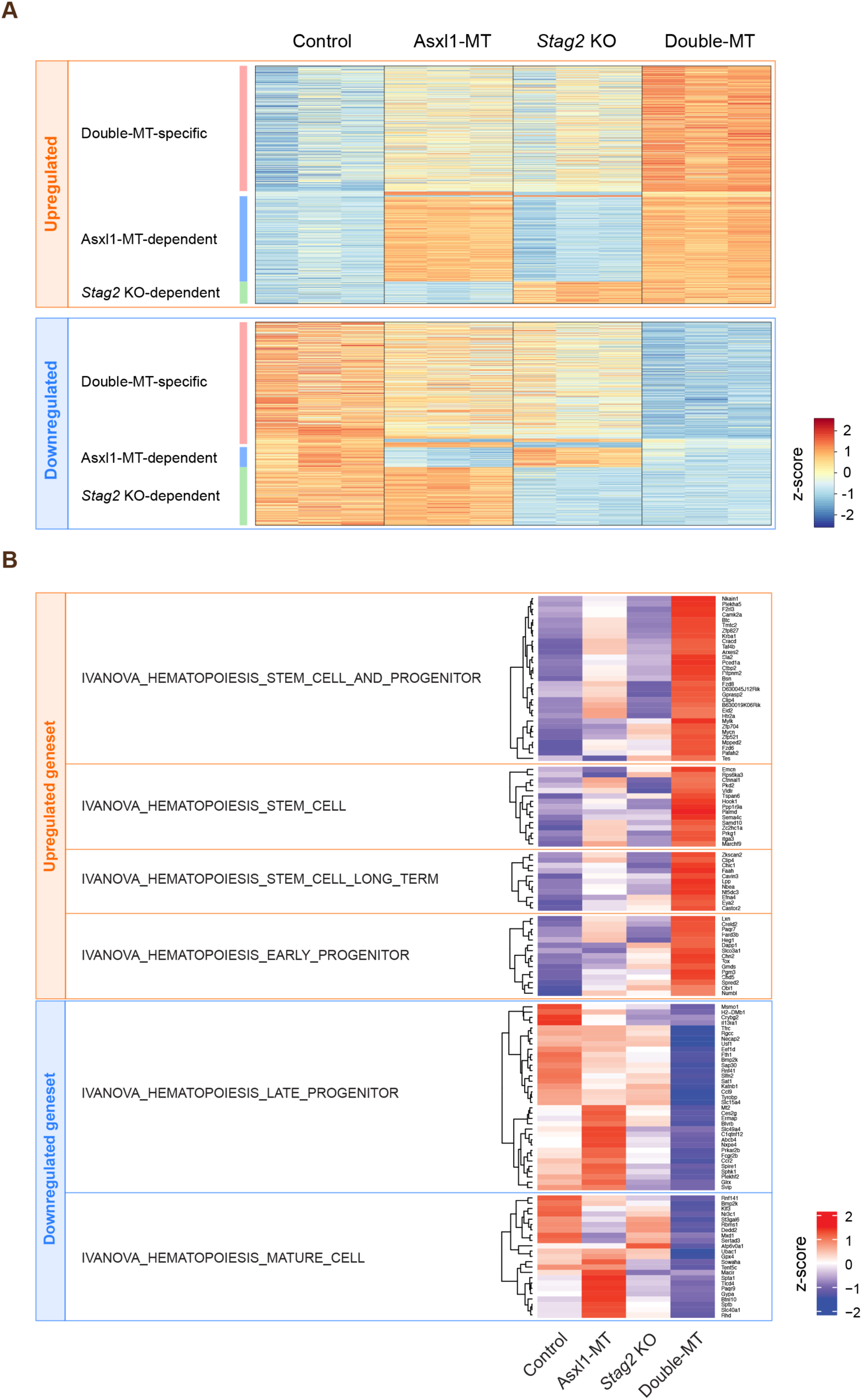
Double-MT mice present specific patterns of gene expressions and histone modifications. **(A)** Differentially expressed genes in double-MT mice are classified into Asxl1-MT-dependent, *Stag2* KO-dependent, and double-MT-specific differentially expressed genes according to expression patterns among genotypes. Expression levels are expressed as z-scores. Double-MT-specific differentially expressed genes are defined as genes whose expression levels are significantly changed in double-MT mice but not in Asxl1-MT KI or *Stag2* KO mice. Asxl1-MT-dependent differentially expressed genes are defined as genes whose expression levels are significantly changed in Asxl1-MT KI and double-MT mice but not in *Stag2* KO mice. *Stag2* KO-dependent differentially expressed genes are defined as genes whose expression levels are significantly changed in *Stag2* KO and double-MT mice but not in Asxl1-MT KI mice. **(B)** Expression levels of genes in significantly upregulated or downregulated CGP (chemical and genetic perturbations) gene sets in the Molecular Signatures Database (MSigDB). Expression levels are expressed as z-scores.

**Figure S5.**
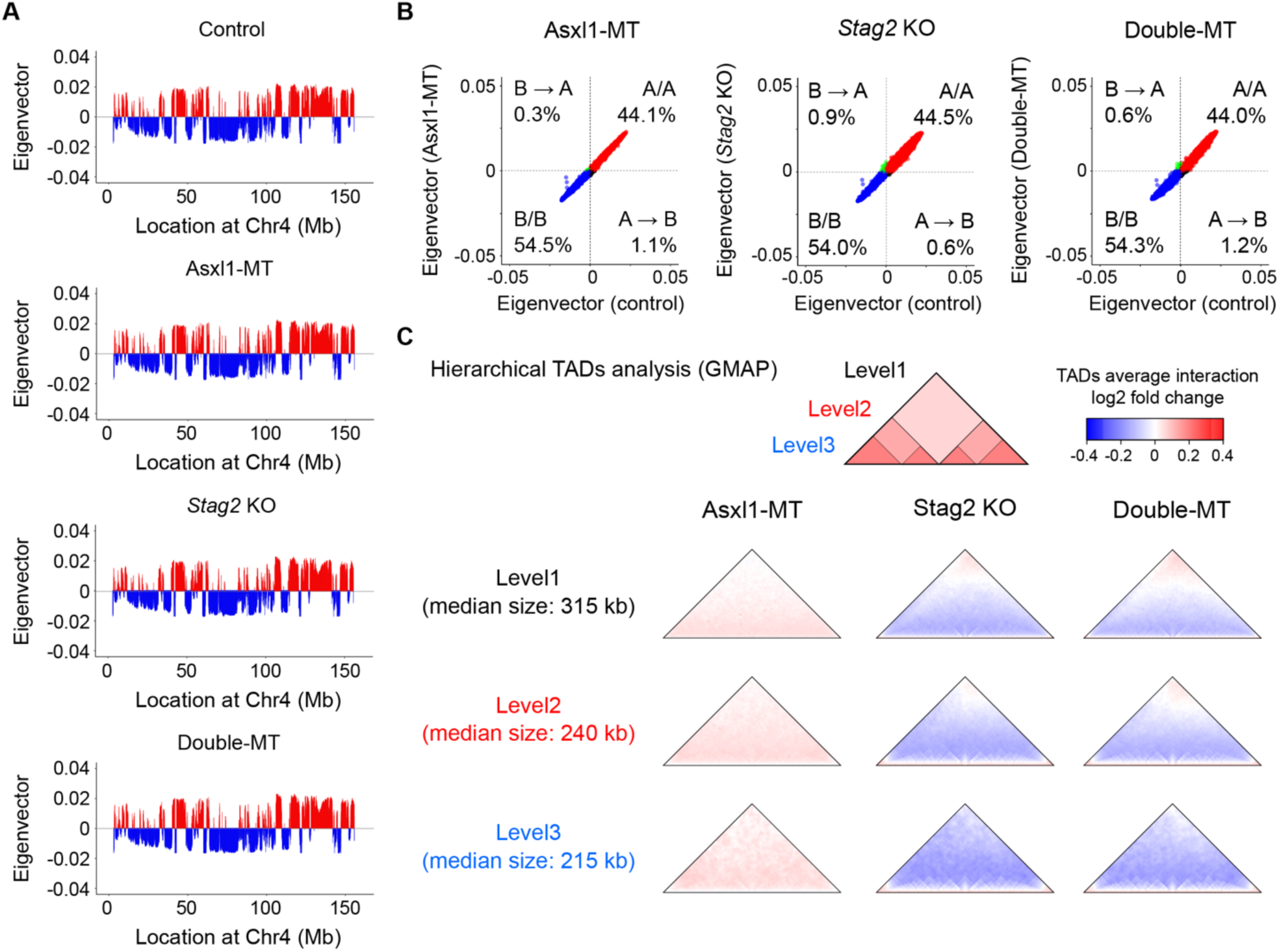
The effects of *Asxl1* and *Stag2* mutations on large-scale genome structures. **(A)** First eigenvalues at each genomic bin on chromosome 4 are shown. Genomic bins with positive and negative eigenvectors indicate A and B compartments, respectively. **(B)** A scatter plot showing the first eigenvalues in each genotype versus control. The frequencies of the bins where assignment to A and B compartments changed between Asxl1-MT KI, *Stag2* KO, or double-MT cells and control cells are noted. The color of the dots represents changed (green: B to A, black: A to B) and unchanged (red: A to A, blue: B to B) bins. **(C)** Differential changes in interactions within each hierarchical level of size-normalized TADs in each genotype compared with control. Hierarchical TADs were called with the Gaussian Mixture model And Proportion test (GMAP), and then each level of TADs shown in the top left panel was separately analyzed.

**Figure S6.**
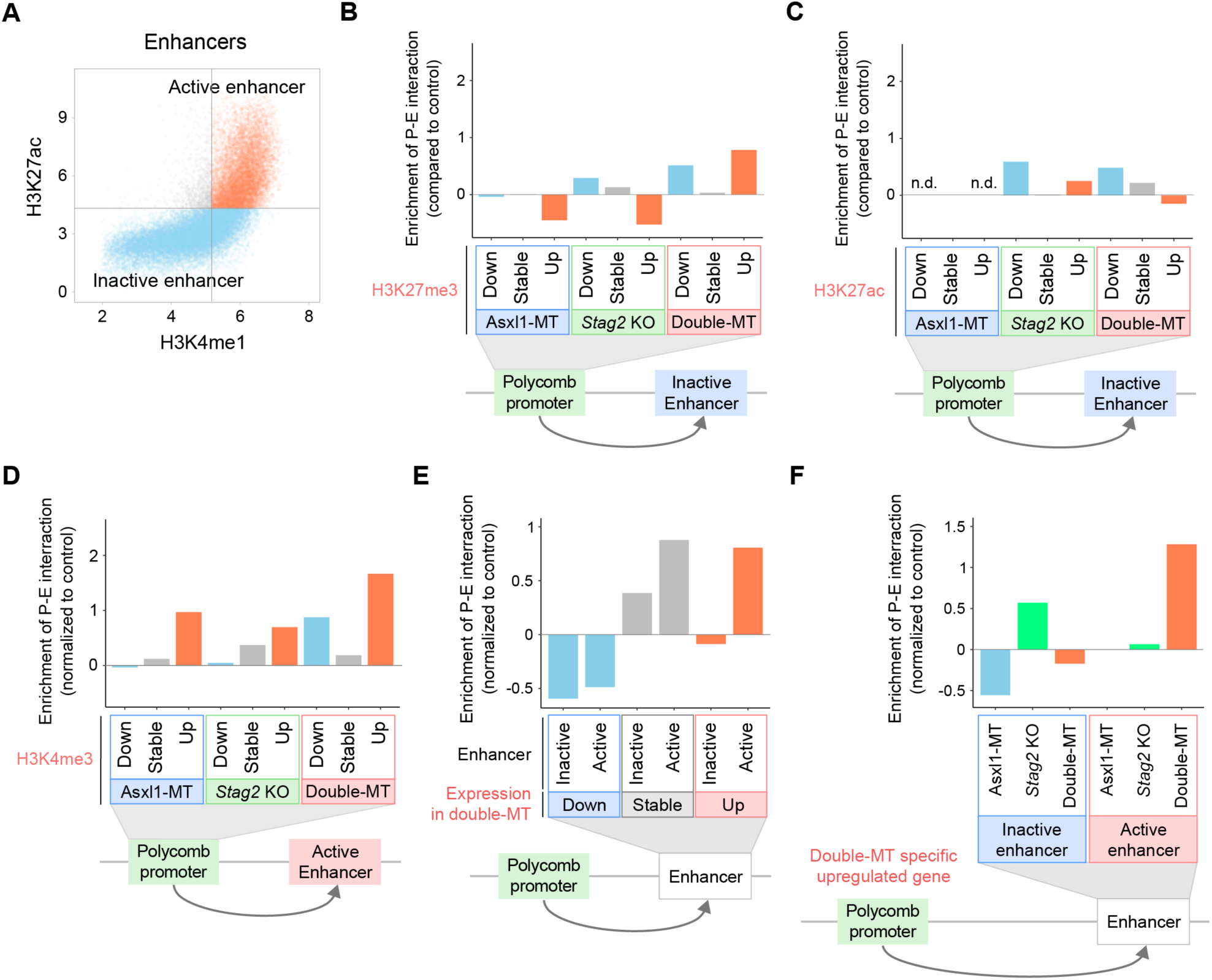
Supplementary data on Hi-C analysis. **(A)** Scatter plots depicting enhancer classifications. Active enhancers are inactive enhancers are defined as H3K4me1^+^/H3K27ac^+^ (orange dot) and H3K27ac^-^ (blue dot) populations, respectively (Creyghton et al., 2010; Lara-Astiaso et al., 2014). **(B, C)** Enrichment of P-E interactions between polycomb-regulated promoters with differential H3K27me3 **(B)** or H3K27ac **(C)** levels (vs. control) and inactive enhancers in each genotype (vs. control). **(D)** Enrichment of P-E interactions between polycomb-regulated promoters with differential H3K4me3 levels (vs. control) and active enhancers in each genotype (vs. control). **(E)** Enrichment of P-E interactions between (i) polycomb-regulated promoters of upregulated, stable, or downregulated genes and (ii) inactive or active enhancers in double-MT cells (vs. control). **(F)** Enrichment of P-E interactions between (i) polycomb-regulated promoters of double-MT-specific upregulated genes and (ii) inactive or active enhancers in each genotype (vs. control). Enrichment is defined as the ratio of observed (obs) number to expected (exp) number. Negative and positive enrichment values represent depletion and enrichment compared with the expected number of interactions, respectively. The difference in the enrichment of P-E interactions (vs. control) was calculated by subtracting the enrichment values of Asxl1-MT KI, *Stag2* KO, or double-MT cells from that of control cells. Negative and positive values represent depletion and enrichment compared with control cells, respectively.

**Figure S7.**
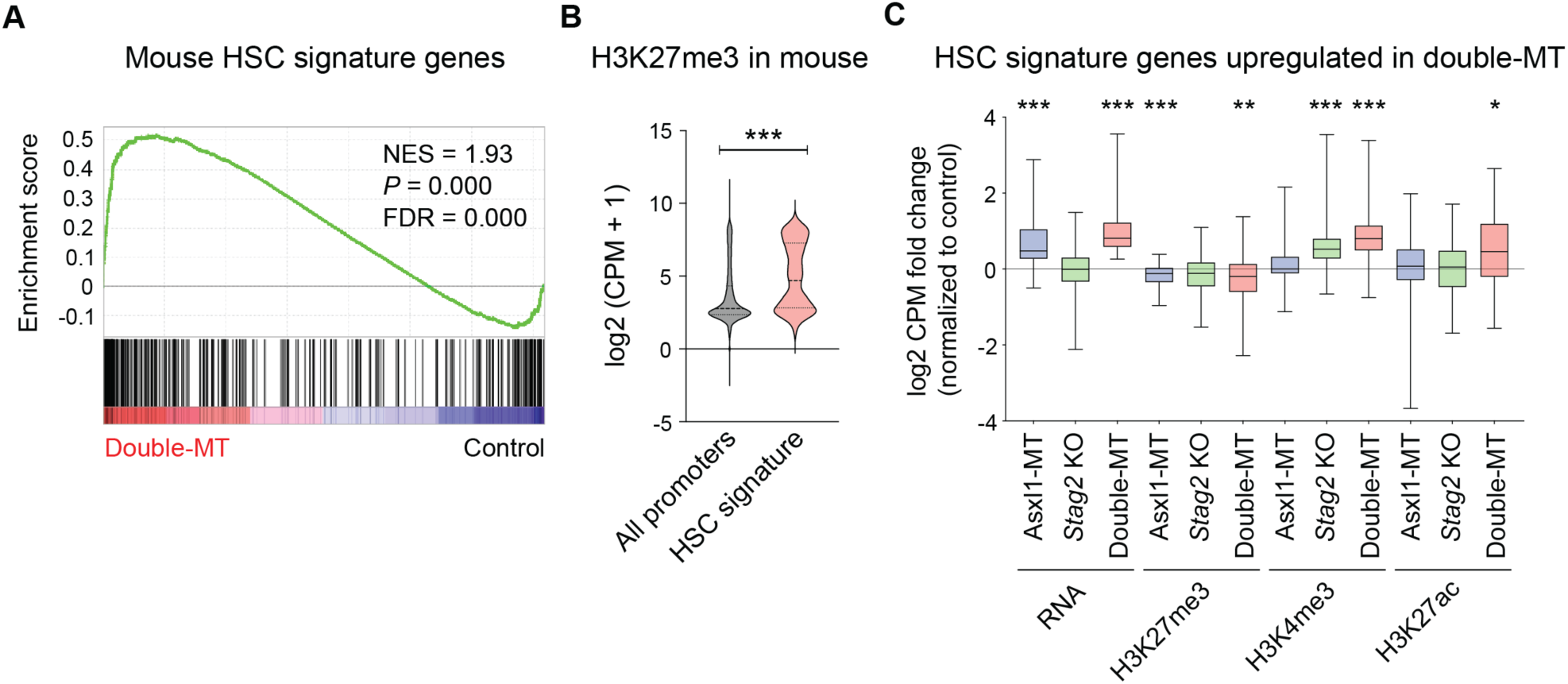
De-repression of polycomb-regulated promoters upregulates genes involved in HSC functions in human and mouse harboring concurrent *ASXL1* and *STAG2* mutations. (A) Gene set enrichment analysis (GSEA) showing the upregulation of HSC signature genes (Chambers et al., 2007) in double-MT mice (vs. control). **(B)** H3K27me3 levels at promoters of mouse HSC signature genes and all pomoters. **(C)** Changes in gene expression and histone modifications levels in HSC signature genes upregulated in double-MT mice. Statistical significance was assessed by two-tailed Student’s t-test **(B)** or one-way ANOVA with Tukey–Kramer’s post-hoc test **(C)**. *P ≤ 0.05, **P ≤ 0.01, ***P ≤ 0.001

## REFERENCES

1. Allis, C.D., and Jenuwein, T. (2016). The molecular hallmarks of epigenetic control. Nat. Rev. Genet. 17, 487–500. 10.1038/nrg.2016.59.

2. Schoenfelder, S., and Fraser, P. (2019). Long-range enhancer–promoter contacts in gene expression control. Nat. Rev. Genet. 10.1038/s41576-019-0128-0.

3. Zheng, H., and Xie, W. (2019). The role of 3D genome organization in development and cell differentiation. Nat. Rev. Mol. Cell Biol. 20, 535–550. 10.1038/s41580-019-0132-4.

4. Stadhouders, R., Filion, G.J., and Graf, T. (2019). Transcription factors and 3D genome conformation in cell-fate decisions. Nature 569, 345–354. 10.1038/s41586-019-1182-7.

5. Lieberman-aiden, E., Berkum, N.L. Van, Williams, L., Imakaev, M., Ragoczy, T., Telling, A., Amit, I., Lajoie, B.R., Sabo, P.J., Dorschner, M.O., et al. (2009). Comprehensive Mapping of Long-Range Interactions Reveals Folding Principles of the Human Genome. Science (80-.). 33292, 289–294.

6. Rao, S.S.P., Huntley, M.H., Durand, N.C., Stamenova, E.K., Bochkov, I.D., Robinson, J.T., Sanborn, A.L., Machol, I., Omer, A.D., Lander, E.S., et al. (2014). A 3D map of the human genome at kilobase resolution reveals principles of chromatin looping. Cell 159, 1665–1680. 10.1016/j.cell.2014.11.021.

7. Dixon, J.R., Selvaraj, S., Yue, F., Kim, A., Li, Y., Shen, Y., Hu, M., Liu, J.S., and Ren, B. (2012). Topological domains in mammalian genomes identified by analysis of chromatin interactions. Nature 485, 376–380. 10.1038/nature11082.

8. Nora, E.P., Lajoie, B.R., Schulz, E.G., Giorgetti, L., Okamoto, I., Servant, N., Piolot, T., Van Berkum, N.L., Meisig, J., Sedat, J., et al. (2012). Spatial partitioning of the regulatory landscape of the X-inactivation centre. Nature 485, 381–385. 10.1038/nature11049.

9. Sanborn, A.L., Rao, S.S.P., Huang, S.C., Durand, N.C., Huntley, M.H., Jewett, A.I., Bochkov, I.D., Chinnappan, D., Cutkosky, A., Li, J., et al. (2015). Chromatin extrusion explains key features of loop and domain formation in wild-type and engineered genomes. Proc. Natl. Acad. Sci. U. S. A. 112, E6456–E6465. 10.1073/pnas.1518552112.

10. Alipour, E., and Marko, J.F. (2012). Self-organization of domain structures by DNA-loop-extruding enzymes. Nucleic Acids Res. 40, 11202–11212. 10.1093/nar/gks925.

11. Fudenberg, G., Imakaev, M., Lu, C., Goloborodko, A., Abdennur, N., and Mirny, L.A. (2016). Formation of Chromosomal Domains by Loop Extrusion. Cell Rep. 15, 2038–2049. 10.1016/j.celrep.2016.04.085.

12. Schwarzer, W., Abdennur, N., Goloborodko, A., Pekowska, A., Fudenberg, G., Loe-Mie, Y., Fonseca, N.A., Huber, W., Haering, C.H., Mirny, L., et al. (2017). Two independent modes of chromatin organization revealed by cohesin removal. Nature 551, 51–56. 10.1038/nature24281.

13. Rao, S.S.P., Huang, S.C., Glenn St Hilaire, B., Engreitz, J.M., Perez, E.M., Kieffer-Kwon, K.R., Sanborn, A.L., Johnstone, S.E., Bascom, G.D., Bochkov, I.D., et al. (2017). Cohesin Loss Eliminates All Loop Domains. Cell 171, 305–320.e24. 10.1016/j.cell.2017.09.026.

14. Dowen, J.M., Fan, Z.P., Hnisz, D., Ren, G., Abraham, B.J., Zhang, L.N., Weintraub, A.S., Schuijers, J., Lee, T.I., Zhao, K., et al. (2014). Control of cell identity genes occurs in insulated neighborhoods in mammalian chromosomes. Cell 159, 374–387. 10.1016/j.cell.2014.09.030.

15. Lupiáñez, D.G., Kraft, K., Heinrich, V., Krawitz, P., Brancati, F., Klopocki, E., Horn, D., Kayserili, H., Opitz, J.M., Laxova, R., et al. (2015). Disruptions of topological chromatin domains cause pathogenic rewiring of gene-enhancer interactions. Cell 161, 1012–1025. 10.1016/j.cell.2015.04.004.

16. Sun, F., Chronis, C., Kronenberg, M., Chen, X.F., Su, T., Lay, F.D., Plath, K., Kurdistani, S.K., and Carey, M.F. (2019). Promoter-Enhancer Communication Occurs Primarily within Insulated Neighborhoods. Mol. Cell 73, 250–263.e5. 10.1016/j.molcel.2018.10.039.

17. Kagey, M.H., Newman, J.J., Bilodeau, S., Zhan, Y., Orlando, D.A., Van Berkum, N.L., Ebmeier, C.C., Goossens, J., Rahl, P.B., Levine, S.S., et al. (2010). Mediator and cohesin connect gene expression and chromatin architecture. Nature 467, 430–435. 10.1038/nature09380.

18. Phillips-Cremins, J.E., Sauria, M.E.G., Sanyal, A., Gerasimova, T.I., Lajoie, B.R., Bell, J.S.K., Ong, C.T., Hookway, T.A., Guo, C., Sun, Y., et al. (2013). Architectural protein subclasses shape 3D organization of genomes during lineage commitment. Cell 153, 1281–1295. 10.1016/j.cell.2013.04.053.

19. Weintraub, A.S., Li, C.H., Zamudio, A. V., Sigova, A.A., Hannett, N.M., Day, D.S., Abraham, B.J., Cohen, M.A., Nabet, B., Buckley, D.L., et al. (2017). YY1 Is a Structural Regulator of Enhancer-Promoter Loops. Cell 171, 1573–1588.e28. 10.1016/j.cell.2017.11.008.

20. Rhodes, J.D.P., Feldmann, A., Hernández-Rodríguez, B., Díaz, N., Brown, J.M., Fursova, N.A., Blackledge, N.P., Prathapan, P., Dobrinic, P., Huseyin, M.K., et al. (2020). Cohesin Disrupts Polycomb-Dependent Chromosome Interactions in Embryonic Stem Cells. Cell Rep. 30, 820–835.e10. 10.1016/j.celrep.2019.12.057.

21. Thiecke, M.J., Wutz, G., Muhar, M., Tang, W., Bevan, S., Malysheva, V., Stocsits, R., Neumann, T., Zuber, J., Fraser, P., et al. (2020). Cohesin-Dependent and-Independent Mechanisms Mediate Chromosomal Contacts between Promoters and Enhancers. Cell Rep. 32, 107929. 10.1016/j.celrep.2020.107929.

22. Kon, A., Shih, L.Y., Minamino, M., Sanada, M., Shiraishi, Y., Nagata, Y., Yoshida, K., Okuno, Y., Bando, M., Nakato, R., et al. (2013). Recurrent mutations in multiple components of the cohesin complex in myeloid neoplasms. Nat. Genet. 45, 1232–1237. 10.1038/ng.2731.

23. Ochi, Y., Kon, A., Sakata, T., Nakagawa, M.M., Nakazawa, N., Kakuta, M., Kataoka, K., Koseki, H., Nakayama, M., Morishita, D., et al. (2020). Combined Cohesin–RUNX1 Deficiency Synergistically Perturbs Chromatin Looping and Causes Myelodysplastic Syndromes. Cancer Discov. 10.1158/2159-8290.cd-19-0982.

24. Tsai, C.H., Hou, H.A., Tang, J.L., Kuo, Y.Y., Chiu, Y.C., Lin, C.C., Liu, C.Y., Tseng, M.H., Lin, T.Y., Liu, M.C., et al. (2017). Prognostic impacts and dynamic changes of cohesin complex gene mutations in de novo acute myeloid leukemia. Blood Cancer J. 7. 10.1038/s41408-017-0022-y.

25. Boultwood, J., Perry, J., Pellagatti, A., Fernandez-Mercado, M., Fernandez-Santamaria, C., Calasanz, M.J., Larrayoz, M.J., Garcia-Delgado, M., Giagounidis, A., Malcovati, L., et al. (2010). Frequent mutation of the polycomb-associated gene ASXL1 in the myelodysplastic syndromes and in acute myeloid leukemia. Leukemia 24, 1062–1065. 10.1038/leu.2010.20.

26. Thol, F., Friesen, I., Damm, F., Yun, H., Weissinger, E.M., Krauter, J., Wagner, K., Chaturvedi, A., Sharma, A., Wichmann, M., et al. (2011). Prognostic significance of ASXL1 mutations in patients with myelodysplastic syndromes. J. Clin. Oncol. 29, 2499–2506. 10.1200/JCO.2010.33.4938.

27. Schnittger, S., Eder, C., Jeromin, S., Alpermann, T., Fasan, A., Grossmann, V., Kohlmann, A., Illig, T., Klopp, N., Wichmann, H.E., et al. (2013). ASXL1 exon 12 mutations are frequent in AML with intermediate risk karyotype and are independently associated with an adverse outcome. Leukemia 27, 82–91. 10.1038/leu.2012.262.

28. Inoue, D., Matsumoto, M., Nagase, R., Saika, M., Fujino, T., Nakayama, K.I., and Kitamura, T. (2016). Truncation mutants of ASXL1 observed in myeloid malignancies are expressed at detectable protein levels. Exp. Hematol. 44, 172–176. 10.1016/j.exphem.2015.11.011.

29. Asada, S., Goyama, S., Inoue, D., Shikata, S., Takeda, R., Fukushima, T., Yonezawa, T., Fujino, T., Hayashi, Y., Kawabata, K.C., et al. (2018). Mutant ASXL1 cooperates with BAP1 to promote myeloid leukaemogenesis. Nat. Commun. 9, 1–18. 10.1038/s41467-018-05085-9.

30. Inoue, D., Kitaura, J., Togami, K., Nishimura, K., Enomoto, Y., Uchida, T., Kagiyama, Y., Kawabata, K.C., Nakahara, F., Izawa, K., et al. (2013). Myelodysplastic syndromes are induced by histone methylation–altering ASXL1 mutations. J Clin Invest. 123, 4627–4640. 10.1172/JCI70739.through.

31. Nagase, R., Inoue, D., Pastore, A., Fujino, T., Hou, H.-A., Yamasaki, N., Goyama, S., Saika, M., Kanai, A., Sera, Y., et al. (2018). Expression of mutant Asxl1 perturbs hematopoiesis and promotes susceptibility to leukemic transformation. J. Exp. Med. 215, 1729–1747. 10.1084/jem.20171151.

32. Surdez, D., Zaidi, S., Grossetête, S., Laud-Duval, K., Ferre, A.S., Mous, L., Vourc’h, T., Tirode, F., Pierron, G., Raynal, V., et al. (2021). STAG2 mutations alter CTCF-anchored loop extrusion, reduce cis-regulatory interactions and EWSR1-FLI1 activity in Ewing sarcoma. Cancer Cell 39, 810–826.e9. 10.1016/j.ccell.2021.04.001.

33. Adane, B., Alexe, G., Seong, B.K.A., Lu, D., Hwang, E.E., Hnisz, D., Lareau, C.A., Ross, L., Lin, S., Dela Cruz, F.S., et al. (2021). STAG2 loss rewires oncogenic and developmental programs to promote metastasis in Ewing sarcoma. Cancer Cell 39, 827–844.e10. 10.1016/j.ccell.2021.05.007.

34. Fujino, T., Goyama, S., Sugiura, Y., Inoue, D., Asada, S., Yamasaki, S., Matsumoto, A., Yamaguchi, K., Isobe, Y., Tsuchiya, A., et al. (2021). Mutant ASXL1 induces age-related expansion of phenotypic hematopoietic stem cells through activation of Akt/mTOR pathway. Nat. Commun. 12, 1–20. 10.1038/s41467-021-22053-y.

35. Viny, A.D., Bowman, R.L., Liu, Y., Lavallée, V.-P., Eisman, S.E., Xiao, W., Durham, B.H., Navitski, A., Park, J., Braunstein, S., et al. (2019). Cohesin Members Stag1 and Stag2 Display Distinct Roles in Chromatin Accessibility and Topological Control of HSC Self-Renewal and Differentiation. Cell Stem Cell, 1–15. 10.1016/j.stem.2019.08.003.

36. Guo, G., Luc, S., Marco, E., Lin, T.W., Peng, C., Kerenyi, M.A., Beyaz, S., Kim, W., Xu, J., Das, P.P., et al. (2013). Mapping cellular hierarchy by single-cell analysis of the cell surface repertoire. Cell Stem Cell 13, 492–505. 10.1016/j.stem.2013.07.017.

37. Ye, M., Zhang, H., Yang, H., Koche, R., Staber, P.B., Cusan, M., Levantini, E., Welner, R.S., Bach, C.S., Zhang, J., et al. (2015). Hematopoietic Differentiation Is Required for Initiation of Acute Myeloid Leukemia. Cell Stem Cell 17, 611–623. 10.1016/j.stem.2015.08.011.

38. Kruse, E.A., Loughran, S.J., Baldwin, T.M., Josefsson, E.C., Ellis, S., Watson, D.K., Nurden, P., Metcalf, D., Hilton, D.J., Alexander, W.S., et al. (2009). Dual requirement for the ETS transcription factors Fli-1 and Erg in hematopoietic stem cells and the megakaryocyte lineage. Proc. Natl. Acad. Sci. U. S. A. 106, 13814–13819. 10.1073/pnas.0906556106.

39. Wilson, N.K., Foster, S.D., Wang, X., Knezevic, K., Schütte, J., Kaimakis, P., Chilarska, P.M., Kinston, S., Ouwehand, W.H., Dzierzak, E., et al. (2010). Combinatorial transcriptional control in blood stem/progenitor cells: genome-wide analysis of ten major transcriptional regulators. Cell Stem Cell 7, 532–544. 10.1016/j.stem.2010.07.016.

40. Loughran, S.J., Kruse, E.A., Hacking, D.F., de Graaf, C.A., Hyland, C.D., Willson, T.A., Henley, K.J., Ellis, S., Voss, A.K., Metcalf, D., et al. (2008). The transcription factor Erg is essential for definitive hematopoiesis and the function of adult hematopoietic stem cells. Nat. Immunol. 9, 810–819. 10.1038/ni.1617.

41. Staber, P.B., Zhang, P., Ye, M., Welner, R.S., Nombela-Arrieta, C., Bach, C., Kerenyi, M., Bartholdy, B.A., Zhang, H., Alberich-Jordà, M., et al. (2013). Sustained PU.1 levels balance cell-cycle regulators to prevent exhaustion of adult hematopoietic stem cells. Mol. Cell 49, 934–946. 10.1016/j.molcel.2013.01.007.

42. Yu, S., Cui, K., Jothi, R., Zhao, D.-M., Jing, X., Zhao, K., and Xue, H.-H. (2011). GABP controls a critical transcription regulatory module that is essential for maintenance and differentiation of hematopoietic stem/progenitor cells. Blood 117, 2166–2178. 10.1182/blood-2010-09-306563.

43. Creyghton, M.P., Cheng, A.W., Welstead, G.G., Kooistra, T., Carey, B.W., Steine, E.J., Hanna, J., Lodato, M.A., Frampton, G.M., Sharp, P.A., et al. (2010). Histone H3K27ac separates active from poised enhancers and predicts developmental state. Proc. Natl. Acad. Sci. U. S. A. 107, 21931–21936. 10.1073/pnas.1016071107.

44. Lara-Astiaso, D., Weiner, A., Lorenzo-Vivas, E., Zaretsky, I., Jaitin, D.A., David, E., Keren-Shaul, H., Mildner, A., Winter, D., Jung, S., et al. (2014). Immunogenetics. Chromatin state dynamics during blood formation. Science 345, 943–949. 10.1126/science.1256271.

45. Schoenfelder, S., Furlan-Magaril, M., Mifsud, B., Tavares-Cadete, F., Sugar, R., Javierre, B.M., Nagano, T., Katsman, Y., Sakthidevi, M., Wingett, S.W., et al. (2015). The pluripotent regulatory circuitry connecting promoters to their long-range interacting elements. Genome Res. 25, 582–597. 10.1101/gr.185272.114.

46. Javierre, B.M., Sewitz, S., Cairns, J., Wingett, S.W., Várnai, C., Thiecke, M.J., Freire-Pritchett, P., Spivakov, M., Fraser, P., Burren, O.S., et al. (2016). Lineage-Specific Genome Architecture Links Enhancers and Non-coding Disease Variants to Target Gene Promoters. Cell 167, 1369–1384.e19. 10.1016/j.cell.2016.09.037.

47. Cairns, J., Freire-Pritchett, P., Wingett, S.W., Várnai, C., Dimond, A., Plagnol, V., Zerbino, D., Schoenfelder, S., Javierre, B.M., Osborne, C., et al. (2016). CHiCAGO: Robust detection of DNA looping interactions in Capture Hi-C data. Genome Biol. 17, 1–17. 10.1186/s13059-016-0992-2.

48. Tyner, J.W., Tognon, C.E., Bottomly, D., Wilmot, B., Kurtz, S.E., Savage, S.L., Long, N., Schultz, A.R., Traer, E., Abel, M., et al. (2018). Functional genomic landscape of acute myeloid leukaemia. Nature 562, 526–531. 10.1038/s41586-018-0623-z.

49. Cheng, T., Rodrigues, N., Shen, H., Yang, Y.G., Dombkowski, D., Sykes, M., and Scadden, D.T. (2000). Hematopoietic stem cell quiescence maintained by p21(cip1/waf1). Science (80-.). 287, 1804–1809. 10.1126/science.287.5459.1804.

50. Beagrie, R.A., Scialdone, A., Schueler, M., Kraemer, D.C.A., Chotalia, M., Xie, S.Q., Barbieri, M., De Santiago, I., Lavitas, L.M., Branco, M.R., et al. (2017). Complex multi-enhancer contacts captured by genome architecture mapping. Nature 543, 519–524. 10.1038/nature21411.

51. Cuadrado, A., Giménez-Llorente, D., Kojic, A., Rodríguez-Corsino, M., Cuartero, Y., Martín-Serrano, G., Gómez-López, G., Marti-Renom, M.A., and Losada, A. (2019). Specific Contributions of Cohesin-SA1 and Cohesin-SA2 to TADs and Polycomb Domains in Embryonic Stem Cells. Cell Rep. 27, 3500–3510.e4. 10.1016/j.celrep.2019.05.078.

52. Goel, V.Y., Huseyin, M.K., and Hansen, A.S. (2023). Region Capture Micro-C reveals coalescence of enhancers and promoters into nested microcompartments. Nat. Genet. 55, 1048–1056. 10.1038/s41588-023-01391-1.

53. Hsieh, T.H.S., Cattoglio, C., Slobodyanyuk, E., Hansen, A.S., Rando, O.J., Tjian, R., and Darzacq, X. (2020). Resolving the 3D Landscape of Transcription-Linked Mammalian Chromatin Folding. Mol. Cell 78, 539–553.e8. 10.1016/j.molcel.2020.03.002.

54. Oudelaar, A.M., Davies, J.O.J., Hanssen, L.L.P., Telenius, J.M., Schwessinger, R., Liu, Y., Brown, J.M., Downes, D.J., Chiariello, A.M., Bianco, S., et al. (2018). Single-allele chromatin interactions identify regulatory hubs in dynamic compartmentalized domains. Nat. Genet. 50, 1744–1751. 10.1038/s41588-018-0253-2.

55. Walter, M.J., Shen, D., Ding, L., Shao, J., Koboldt, D.C., Chen, K., Larson, D.E., McLellan, M.D., Dooling, D., Abbott, R., et al. (2012). Clonal Architecture of Secondary Acute Myeloid Leukemia. N. Engl. J. Med. 366, 1090–1098. 10.1056/NEJMoa1106968.

56. Lindsley, R.C., Mar, B.G., Mazzola, E., Grauman, P. V., Shareef, S., Allen, S.L., Pigneux, A., Wetzler, M., Stuart, R.K., Erba, H.P., et al. (2015). Acute myeloid leukemia ontogeny is defined by distinct somatic mutations. Blood 125, 1367–1376. 10.1182/blood-2014-11-610543.

57. Haferlach, T., Nagata, Y., Grossmann, V., Okuno, Y., Bacher, U., Nagae, G., Schnittger, S., Sanada, M., Kon, A., Alpermann, T., et al. (2014). Landscape of genetic lesions in 944 patients with myelodysplastic syndromes. Leukemia 28, 241–247. 10.1038/leu.2013.336.

58. Papaemmanuil, E., Gerstung, M., Malcovati, L., Tauro, S., Gundem, G., Loo, P. Van, Yoon, C.J., Ellis, P., Wedge, D.C., Pellagatti, A., et al. (2013). Clinical and biological implications of driver mutations in myelodysplastic syndromes. Blood 122, 3616–3627. 10.1182/blood-2013-08-518886.The.

59. Makishima, H., Yoshizato, T., Yoshida, K., Sekeres, M.A., Radivoyevitch, T., Suzuki, H., Przychodzen, B.J., Nagata, Y., Meggendorfer, M., Sanada, M., et al. (2017). Dynamics of clonal evolution in myelodysplastic syndromes. Nat. Genet. 49, 204–212. 10.1038/ng.3742.

60. Yoshizato, T., Nannya, Y., Atsuta, Y., Shiozawa, Y., Iijima-Yamashita, Y., Yoshida, K., Shiraishi, Y., Suzuki, H., Nagata, Y., Sato, Y., et al. (2017). Genetic abnormalities in myelodysplasia and secondary acute myeloid leukemia: impact on outcome of stem cell transplantation. Blood 129, 2347–2358. 10.1182/blood-2016-12-754796.

61. Meggendorfer, M., de Albuquerque, A., Nadarajah, N., Alpermann, T., Kern, W., Steuer, K., Perglerova, K., Haferlach, C., Schnittger, S., and Haferlach, T. (2015). Karyotype evolution and acquisition of FLT3 or RAS pathway alterations drive progression of myelodysplastic syndrome to acute myeloid leukemia. Haematologica 100, e487–e490. 10.3324/haematol.2015.127985.

62. Benjamini, Y., and Hochberg, Y. (1995). Controlling the False Discovery Rate: A Practical and Powerful Approach to Multiple Testing. J. R. Stat. Soc. Ser. B Stat. Methodol. 57, 289–300. 10.1111/j.2517-6161.1995.tb02031.x.

63. Heinz, S., Benner, C., Spann, N., Bertolino, E., Lin, Y.C., Laslo, P., Cheng, J.X., Murre, C., Singh, H., and Glass, C.K. (2010). Simple combinations of lineage-determining transcription factors prime cis-regulatory elements required for macrophage and B cell identities. Mol. Cell 38, 576–589. 10.1016/j.molcel.2010.05.004.

64. Robinson, M.D., McCarthy, D.J., and Smyth, G.K. (2010). edgeR: a Bioconductor package for differential expression analysis of digital gene expression data. Bioinformatics 26, 139–140. 10.1093/bioinformatics/btp616.

65. Zhang, Y., Liu, T., Meyer, C.A., Eeckhoute, J., Johnson, D.S., Bernstein, B.E., Nusbaum, C., Myers, R.M., Brown, M., Li, W., et al. (2008). Model-based Analysis of ChIP-Seq (MACS). Genome Biol. 9, R137. 10.1186/gb-2008-9-9-r137.

66. Gu, Z., Eils, R., and Schlesner, M. (2016). Complex heatmaps reveal patterns and correlations in multidimensional genomic data. Bioinformatics 32, 2847–2849. 10.1093/bioinformatics/btw313.

67. Tibshirani, R., Walther, G., and Hastie, T. (2001). Estimating the Number of Clusters in a Data Set Via the Gap Statistic. J. R. Stat. Soc. Ser. B Stat. Methodol. 63, 411–423. 10.1111/1467-9868.00293.

68. Roayaei Ardakany, A., Gezer, H.T., Lonardi, S., and Ay, F. (2020). Mustache: multi-scale detection of chromatin loops from Hi-C and Micro-C maps using scale-space representation. Genome Biol. 21, 256. 10.1186/s13059-020-02167-0.

69. Yu, W., He, B., and Tan, K. (2017). Identifying topologically associating domains and subdomains by Gaussian Mixture model And Proportion test. Nat. Commun. 8, 535. 10.1038/s41467-017-00478-8.

70. Ramírez, F., Bhardwaj, V., Arrigoni, L., Lam, K.C., Grüning, B.A., Villaveces, J., Habermann, B., Akhtar, A., and Manke, T. (2018). High-resolution TADs reveal DNA sequences underlying genome organization in flies. Nat. Commun. 9, 189. 10.1038/s41467-017-02525-w.

71. Harmston, N., Ing-Simmons, E., Perry, M., Barešić, A., and Lenhard, B. (2015). GenomicInteractions: An R/Bioconductor package for manipulating and investigating chromatin interaction data. BMC Genomics 16, 963. 10.1186/s12864-015-2140-x.

72. Jaatinen, T., Hemmoranta, H., Hautaniemi, S., Niemi, J., Nicorici, D., Laine, J., Yli-Harja, O., and Partanen, J. (2006). Global Gene Expression Profile of Human Cord Blood–Derived CD133+ Cells. Stem Cells 24, 631–641. 10.1634/stemcells.2005-0185.

73. Chambers, S.M., Boles, N.C., Lin, K.Y.K., Tierney, M.P., Bowman, T. V., Bradfute, S.B., Chen, A.J., Merchant, A.A., Sirin, O., Weksberg, D.C., et al. (2007). Hematopoietic Fingerprints: An Expression Database of Stem Cells and Their Progeny. Cell Stem Cell 1, 578–591. 10.1016/j.stem.2007.10.003.

